# Intergenerational instability of the C9orf72 hexanucleotide repeat

**DOI:** 10.64898/2026.09.10.750611

**Authors:** Osma S. Rautila, Anna Kiviharju, Lilja Jansson, FinnGen, Karri Kaivola, Pentti J. Tienari

**Affiliations:** Translational Immunology, Research Programs Unit, University of Helsinki, Helsinki, Finland; Department of Neurology, Helsinki University Hospital, Helsinki, Finland; Clinical Genetics Unit, HUS Diagnostic Center, Helsinki University Hospital, Helsinki, Finland

**Author notes:** Equal contribution (Karri Kaivola, Pentti J. Tienari). Corresponding author:* Osma S. Rautila.

**Keywords:** repeat disorders, repeat instability, parental transmission, C9orf72, GGGGCC

## Abstract

The *C9orf72* hexanucleotide repeat expansion (HRE) is the most common cause of amyotrophic lateral sclerosis (ALS) and frontotemporal dementia (FTD). It follows autosomal dominant inheritance in families; however, a high proportion of cases are sporadic, raising the possibility of parental premutation. We have demon strated that intermediatelength alleles (IAs) with ≥ 18 repeats (allele frequency ≈1%) belong to the same pool of haplotypes as the HRE, suggesting shared ancestry. Here, we tested whether alleles with ≥ 18 repeats expand in parental transmission. We used two repeat-primed PCR methods to analyze allele lengths in 539 genetically unselected parent-offspring pairs and in 152 pairs known to carry the SNP (rs139185008*C) that tags ≥ 18–20 repeat IAs and the HRE in Finland. We discovered intergenerational repeat length changes only in ≥ 20 repeat alleles. A significant (P = 0.0059) sex bias in 6–40 repeat alleles was noted using a logistic regression model. In this allele range, 12 out of 16 expansions were paternally inherited and 6 out of 7 contractions were maternally inherited. The expansion rate of 20–40 repeat alleles was 34 % in paternal and 11 % in maternal transmissions. In the 20–40 repeat range, most intergenerational expansions were 1–4 repeats in size (15/16), but one larger “jump”, a paternal expansion from 27 to 73 repeats, was observed. These results demonstrate that alleles with ≥ 20 repeats have an increased likelihood of instability, that a paternal expansion bias is observed in alleles with 20–40 repeats, and that expansion events are predominantly 1–4 repeats in size.

## INTRODUCTION

The *C9orf72* hexanucleotide repeat expansion (HRE) in the first intron of *C9orf72* is a major cause of amyotrophic lateral sclerosis (ALS) and frontotemporal dementia (FTD).^**1–4**^ Despite autosomal dominant inheritance, many *C9orf72*-related ALS patients have no family history of ALS. One hypothesis for the common occurrence of sporadic *C9orf72*-related ALS is that an unstable, expansion-prone premutation could be present in one of the parents. Many repeat expansion disorders exhibit both intergenerational and somatic instability.^**5**^ Certain patterns of instability are also shared between disorders. For example, a tendency for repeat tract expansions exists during paternal transmission in Huntington’s disease (caused by CAG repeats),^**6**^ while in Friedreich’s ataxia (GAA) and spinocerebellar ataxia type 27b (GAA), expansion during maternal transmission is reportedly more likely.^**7,8**^ The parental sex bias can be repeat-length dependent: in a study of 5508 transmissions in Fragile X (CGG), short premutation alleles had a higher rate of expansion paternally, but longer alleles maternally. Also, as the repeat units increased, more contractions were seen in paternal, but not in maternal transmission.^**9**^

For the *C9orf72* G4C2 repeat, most studies on intergenerational instability have focused on either expansion carriers or short repeat sizes. A dataset of 1046 parent-offspring pairs from lymphoblastoid cell line DNA contained two paternal expansions and one paternal contraction: from 21 repeats to 22, from 22 to 20, and from 11 to 12. The alleles with less than 10 repeats were stable.^**10**^ As the cohort selection criteria were not related to *C9orf72* G4C2, only a few large intermediate alleles were present, and the largest parental allele was just 22 repeats. However, it is only the larger alleles with ~70 and 120 repeat units that have been reported to undergo extensive intergenerational increases in size to HREs in ALS or FTD families.^**11,12**^

In contrast, in HREs, a contraction bias has been reported. In a cohort of 17 parent-offspring pairs with expansions, ten contracted, five remained stable and two expanded. A slight paternal contraction bias was reported and a single very large paternal contraction from 1700 repeats to 100 repeats was noted. In another cohort of 16 parent-offspring pairs, 11 expansions contracted, 1 remained stable and 4 expanded. No sex bias was reported and all inherited HREs that contracted remained over 1000 repeats long.^**13**^

Although direct evidence is lacking for large and small intergenerational expansion events, indirect genetic evidence does exist. In Europe, a founder haplotype tagged by rs3849942*A is shared by both expansions and intermediate-length alleles (IAs) containing ≥7 repeat units^**3,10,14–17**^ and in Finland, a sub-founder haplotype tagged by rs139185008*C is shared by both expansions and even longer IAs often containing ≥18–20 repeat units.^**17**^ Using haplotype-based analyses, we have previously shown that IAs with ≥18–20 repeats belong to the same pool of haplotypes as the HRE. This suggests that the HRE could evolve via repeat instability from alleles with ≥18–20 repeats.^**18**^

Here, we analyzed *C9orf72* G4C2 IA intergenerational instability and the hypothetical role of IAs as premutations. To achieve this, we first analyzed 539 parent-offspring transmissions of C9orf72 G4C2 alleles without genetic selection criteria and then 152 parent-offspring pairs in which both parent and offspring carried the rs139185008*C variant tagging the longer IAs.

## METHODS

As part of previous multiple sclerosis (MS) genetics studies, we genotyped Finnish MS nuclear families with at least one parent available using repeat-primed PCR (RP-PCR).^**19**^ Peripheral blood-derived DNA was available in 460 families.

To enrich for longer IAs and the *C9orf72* HRE, we selected 311 trios from FinnGen,^**20**^ where a child and only one of the parents carried rs139185008*C. As some of the parents had multiple children, this resulted in 169 parent-offspring pairs. The parent-offspring status was confirmed using genome-wide single nucleotide variant genotyping data provided by the biobanks. The genotyping had been performed with the FinnGen ThermoFisher Axiom custom array. For per-sample quality control, we used discordant sex information, > 3 SD heterozygosity rate and > 5 % missing genotype rate as exclusion criteria. Per-variant inclusion criteria included a genotyping rate of ≥ 95 %, Hardy-Weinberg equilibrium (p>0.000001) and minor allele frequency ≥ 0.01. The proportion IBD was calculated with plink v.2.0.0^**21**^ and parent-offspring status was defined with a proportion IBD of 0.45-0.55, Z1 > 0.9 and a ≥ 20-year age gap.

We collected DNA from peripheral blood leukocytes and performed repeat-primed PCR for all samples to assess *C9orf72* G4C2 repeat lengths.^**22**^ Additionally, in the cohort enriched for IAs based on rs139185008*C carriership, all samples with expansions and/or indications of intergenerational instability in standard RP-PCR were reanalyzed with the commercial AmplideX C9orf72 kit (Asuragen, USA). For the few conflicting estimates between standard RP-PCR and the Amplidex C9orf72 kit, the repeat lengths from the Amplidex C9orf72 kit were used. The results were visualized using GeneMapper software 6 (ThermoFisher) and the AmplideX PCR CE C9orf72 Analysis Module v1.0. The longer allele was assumed to be in cis with rs139185008*C.

To analyze the effects of repeat length, parental age, parental sex and offspring sex on intergenerational expansion events, we used a logistic regression model in Python 3.11 with statsmodels v0.14.^**23**^

## RESULTS

In the MS cohort where samples were not selected based on any *C9orf72* haplotype, we successfully genotyped 1081 (88 % success rate) of the 1235 samples after a single round of genotyping. We analyzed 539 parent-offspring transmissions (350 maternal, 189 paternal) from families with at least one parent available. We recorded 526 transmissions of alleles of <18 repeats and detected no expansions or contractions. Eleven transmissions of alleles with 18–31 repeats were recorded (7 maternal, 4 paternal). One suspected expansion (from 23 to 25 repeats) and one contraction (from 22 to 20 repeats) were detected, both from paternal transmissions (Figure S1). The C9orf72 HRE was found in two families, and it was transmitted from the father to the daughter in both families.

Next, we focused on the longer IAs by selecting parent-offspring pairs from two Finnish biobanks (THL and Helsinki Biobank) carrying rs139185008*C, which tags IAs with ≥ 18 repeats and the HRE.^**17**^ DNA was available for 300 samples and we successfully determined the C9orf72 repeat lengths in 284/300 (94.7 %) samples after two rounds of genotyping, representing 152 transmissions between 118 parents and 152 children (Figure 1B).

**Figure 1:**
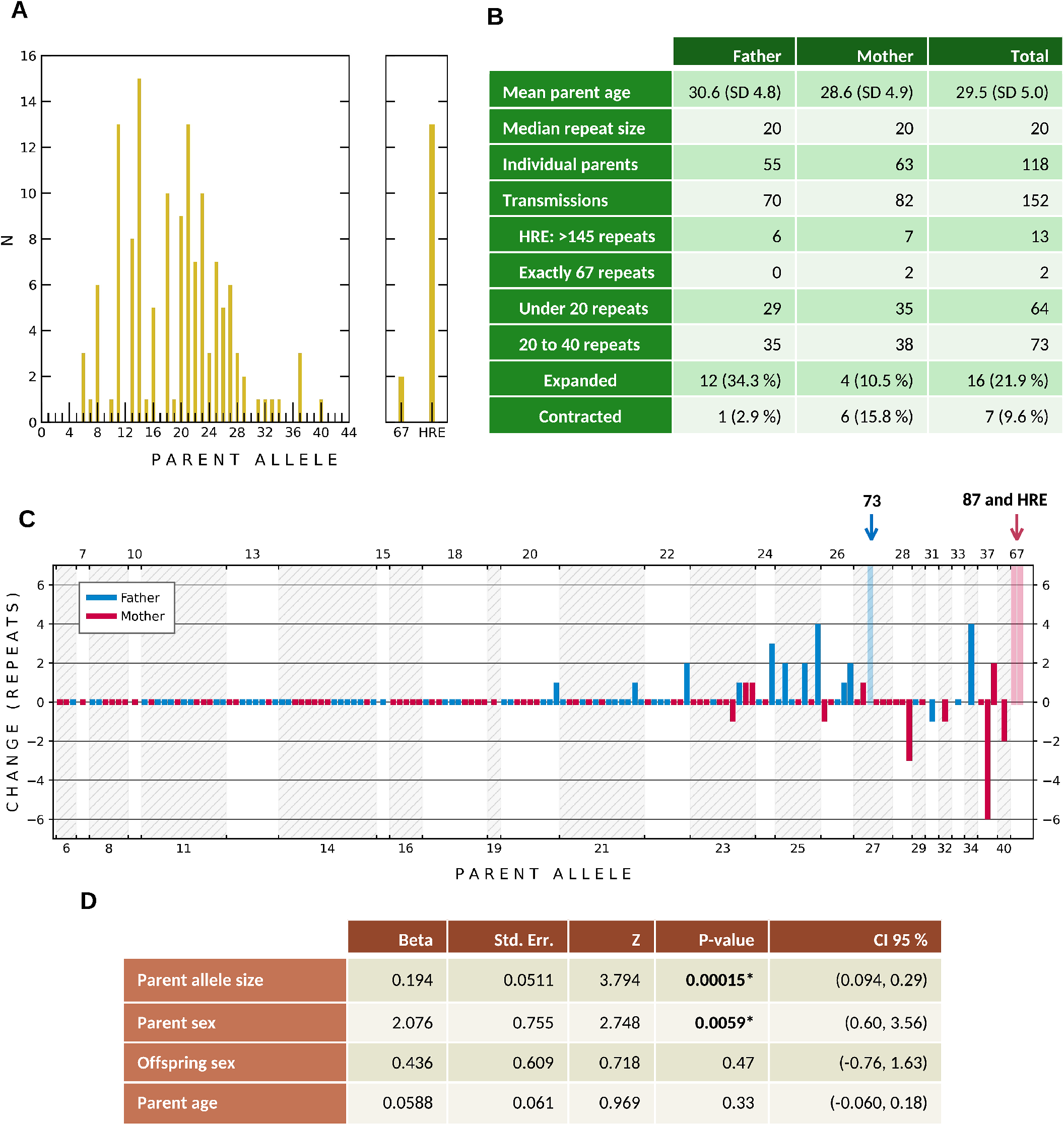
Intergenerational repeat instability. **A**. C9orf72 G4C2 lengths of the longer allele in parental transmissions in biobank data **B**. Summary statistics of biobank data. **C**. Instability in parental transmission in 6 to 40 repeat units by allele size and sex in biobank data. Allele size on the xaxis, change in parental transmission on the yaxis. Paternal transmission in blue, maternal in red. Lower opacity bars with arrows on top denote the transmissions with larger jumps to ~73, ~87 and HRE. **D**. Logistic regression model of expansion events in biobank data. Allele size and paternal transmission are associated with expansions. *Statistically significant (p > 0.05).

The parental allele sizes in the transmissions is shown in Figure 1A. The concordance between in-house RP-PCR and Amplidex was high; however, a few of the longer >35 IAs (n = 9) were misclassified as expansions in the in-house RP-PCR (Table S1). We did not perform over-the-repeat PCR^**22**^ to confirm biallelic amplicons in long IAs in order to save DNA for Amplidex. All transmissions of IAs with < 20 repeats (n=64) were stable. In the transmissions of IAs with 20–40 repeats (n=73), 16 expanded, 7 contracted and 61 remained the same size (Figure 1B and 1C). Most expansions and contractions were in the range of 1 to 4 units. One larger expansion was found in the transmissions of IAs with 20–40 repeats, which was a paternally inherited expansion from 27 repeats to 73 repeats (parent age 42). One mother had two instability events in three transmissions: her allele with 37 repeats was stable once, expanded once to 39 and contracted once to 31 repeats.

There was a marked sex bias in the expansion vs. contraction events in the IAs with 20–40 repeats. Of the expansions 12 of 16 (75 %) were paternally inherited, whereas 6 of 7 (86 %) of the contractions were maternally inherited (Figure 1B and 1C). The expansion rate of 20–40 repeats was 34 % when paternally and 11 % when maternally inherited.

Longer repeats were more unstable: 7 of 13 (54 %) paternal alleles with 25–34 repeats expanded, while only 5 of 23 (22 %) alleles with 20–24 repeat units expanded.

In a logistic regression analysis of the transmissions of alleles with 6–40 repeats, transmission from the father (p = 0.0059) and repeat tract length (p = 0.00015) were significantly associated with an expansion event (Figure 1D).

Outside the IA range of 20–40 repeats, we discovered a mother with ~67 repeats, who transmitted an ~87 repeat allele and the HRE to her two children when she was 29 and 31 years old (Figure 1C). All instability events observed in RP-PCR are shown in Table S1 and Figure S2.

## DISCUSSION

Our results shed light on why apparently sporadic C9orf72-related ALS is so common. In addition to reduced penetrance,^**24,25**^ unstable premutations expanding into pathogenic repeat lengths across generations can be a contributing factor.

In the analysis of parent-offspring transmissions, we found that IAs with 20–40 repeats are unstable in a significant proportion of transmissions. In paternal transmissions 34% of alleles expanded and 3% contracted, while in maternal transmissions 11% expanded and 16% contracted. Most expansion events were just 1–4 repeats in size.

Our data indicate a bi-threshold effect where the likelihood of instability first increases after the repeat length reaches ≥ 20 units. This first threshold is in line with haplotype data where increased haplotype sharing was detected between IAs with ≥ 18 repeats and the HRE,^**18**^ and in vitro data from G4C2 repeats in transfected cells using the SV40 replication system.^**26**^

The second threshold is where the repeat “jumps” to full HRE. This threshold cannot be precisely defined yet. Intergenerational jumps to expansions have been previously reported at ~70 and ~120 repeat units.^**11,12**^ In our data, one mother with ~67 repeats transmitted ~87 repeats and the HRE to her two children. These data suggest that ~67 repeats are already above the second threshold. The length change from 27 repeats to ~73 repeats also indicates that jumps of smaller magnitude can occur at lower repeat sizes. In our data, there were only two alleles in the 40–145 repeat range (Figure 1A), yet 13 HREs were present. Perhaps this dip in allele frequencies indicates that alleles in this range have a high tendency for expansion.

In addition to simple size-dependent instability, our study indicates a paternal bias for expansion in the 20–40 repeat range (p = 0.0059). This does not mean that the full-length expansions are more likely to be of paternal origin. For example, in Fragile X, a parental expansion bias has been shown to reverse to contractions with longer allele sizes of ≥ 95 CGG repeats.^**9**^ Extending the parental bias of Fragile X to the C9orf72 HRE would mean that full-length expansions might be more prone to contraction in paternal transmissions.

The small changes (perhaps 1–6 units) in repeat sizes could be driven by DNA polymerase slippage, but this cannot explain “jumps” from IAs to expansions requiring several multiplications of the repeat tract. Such jumps could either be the result of DNA repair pathways that can produce large expansions (e.g. nick-mediated repeat instability)^**27**^ or multiple failed repair events leading to incremental changes in the repeat tract during mitotic DNA replication and/or post-mitotic transcription-coupled repair.^**28–33**^ The latter implies that somatic instability in gametes would contribute to the jumps. In mice with humanized *C9orf72* alleles,^**34**^ rather large G4C2 repeats (95–300 units) have been, however, reported to be relatively stable, indicating notable differences between these mice and humans. Age could be one factor.

Somatic mosaicism is a sign of somatic repeat instability. In human blood, mosaicism of the C9orf72 G4C2 tract has been reported at ~50–200 repeats,^**12,35–37**^ while 20–27 repeat alleles did not smear in Southern blots.^**37**^ This range coincides with the range where intergenerational jumps to expansion have been reported. However, instability in gametes has not been directly studied, and the tendency for C9orf72 G4C2 to somatically expand is cell type-specific. The human frontal cortex often contains large repeats,^**38**^ whereas, for example, cerebellar cells and fibroblasts contain shorter repeats.^**37–39**^ This is highlighted by reports of subjects with under 100 repeats in blood having over 1000 repeats in the brain. In one report using Southern blotting, a subject had ~70–120 repeats in blood DNA, but over 1000 repeats in the frontal cortex.^**37**^ In a recent long-read sequencing study, a large somatic expansion from a 4-repeat allele was reported in the prefrontal cortex of an FTD patient.^**40**^ This highlights that any proposed thresholds for repeat instability are not absolute, but based on likelihoods. In the long-read sequencing study, a total of four somatic expansion events were detected in ALS and FTD brains with repeat tract lengths starting from 4, 20, 56 and 64 units.^**40**^ This phenomen of IAs expanding to HREs in the brain could hypothetically also explain the increased risk of ALS in IA carriers.^**15,41–44**^ On a related note, the C9orf72 G4C2 repeat has been reported to behave as a folate-sensitive fragile site, and somatic repeat expansion could exacerbate this fragility.^**45**^

Our study has several limitations. First, RP-PCR can only identify repeat sizes up to ~145 repeat units. Second, the interpretation of repeats at > 40 in RP-PCR is challenging owing to both technical limitations and possible somatic mosaicism. Third, a ± 1 repeat unit margin of error exists in RP-PCR. To combat this, our study used two independent genotyping runs. Also, many of the observed changes in size were larger than one repeat. Fourth, the haplotype background was homogeneous, which may affect the results since Finland has the highest reported frequency of *C9orf72* HRE in ALS/FTD. We cannot rule out the possibility that the haplotype tagged by rs139185008*C affects intergenerational instability of alleles with ≥ 20 repeats. Fifth, all DNA samples were extracted from blood and we could not analyze possible tissue-specific effects. Sixth, the study was underpowered for the analysis of genetic modifiers of repeat instability, which represents an important area of future research since C9orf72 G4C2 instability has been reported to depend on the mismatch repair pathway protein Msh2 in C9orf72 G4C2 knock-in mice.^**34**^

## Abbreviations

IA: intermediatelength allele
HRE: hexanucleotide repeat expansion
ALS: amyotrophic lateral sclerosis
FTD: frontotemporal dementia

## CODE AND DATA AVAILABILITY

All raw RP-PCR data are available upon reasonable request.

## ACKNOWLEDGEMENTS

Thank you to FinnGen (access to genotypes for exploration), THL Biobank (reference panel and samples) and Helsinki Biobank (samples). We thank all study participants for their generous participation in the Helsinki and THL Biobanks. Thank you to the FinnGen committee for accepting our study design and the staff at THL and Helsinki Biobanks for their generous help in providing the samples and genotyping data. We also thank CSC for valuable computing resources.

This study was funded by the Finnish Cultural Foundation, the Sigrid Juselius Foundation, the Maire Taponen Foundation, the Finnish Brain Foundation, the Finnish Medical Foundation, the Biomedicum Helsinki Foundation, Helsinki University Hospital grants, the Finnish Academy and the Orion Research Foundation.

## AUTHOR CONTRIBUTIONS

O.S.R., K.K and P.J.T conceptualized and designed research; L.J and A.K performed RP-PCR genotyping; O.S.R analyzed data; O.S.R wrote the manuscript; K.K and P.J.T supervised the work; All authors edited the manuscript.

