## Supplemental Figures for "Intergenerational instability of the C9orf72 hexanucleotide repeat"

**Supplemental Figure 1.** Instability events in the MS cohort

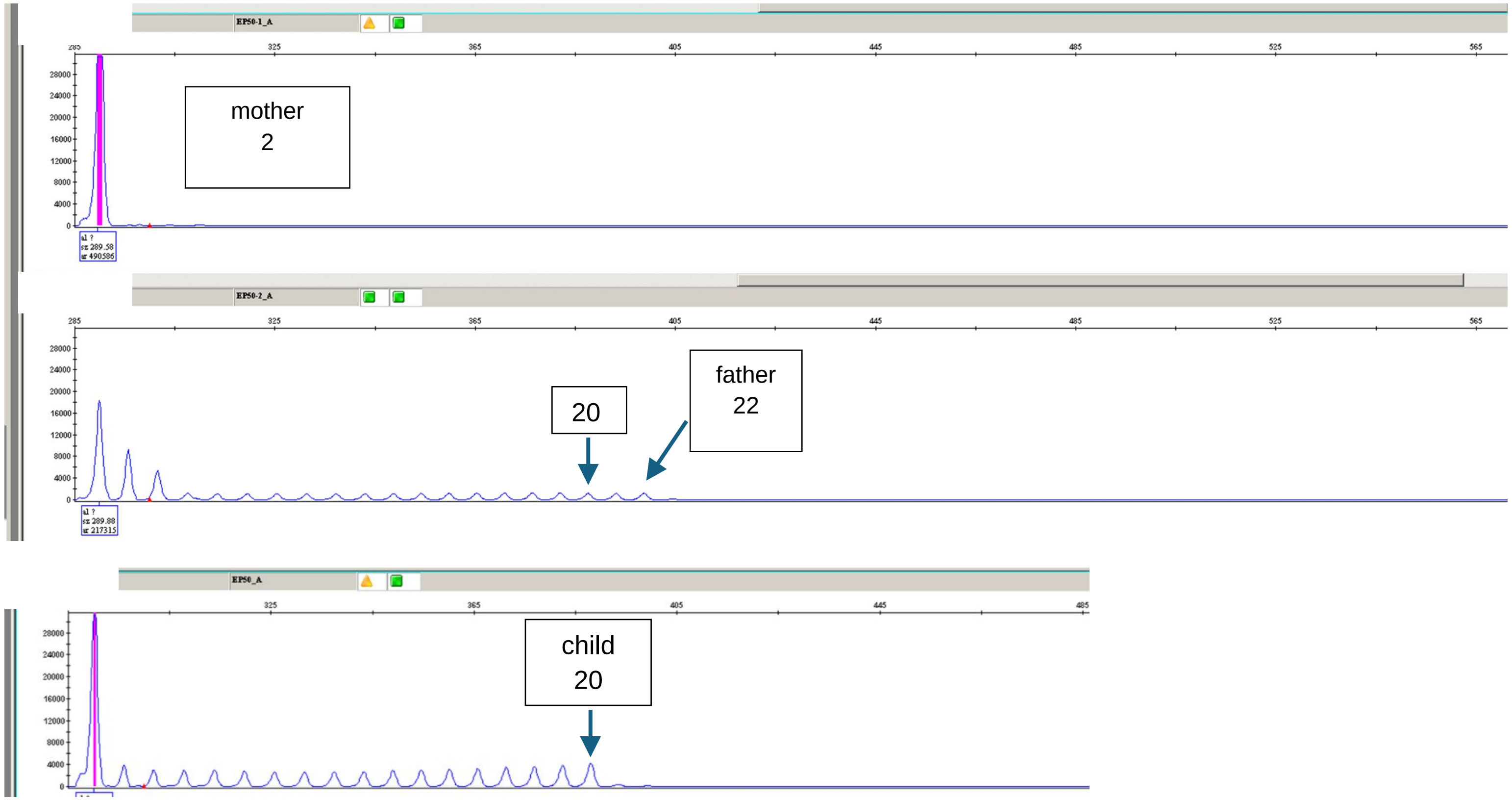

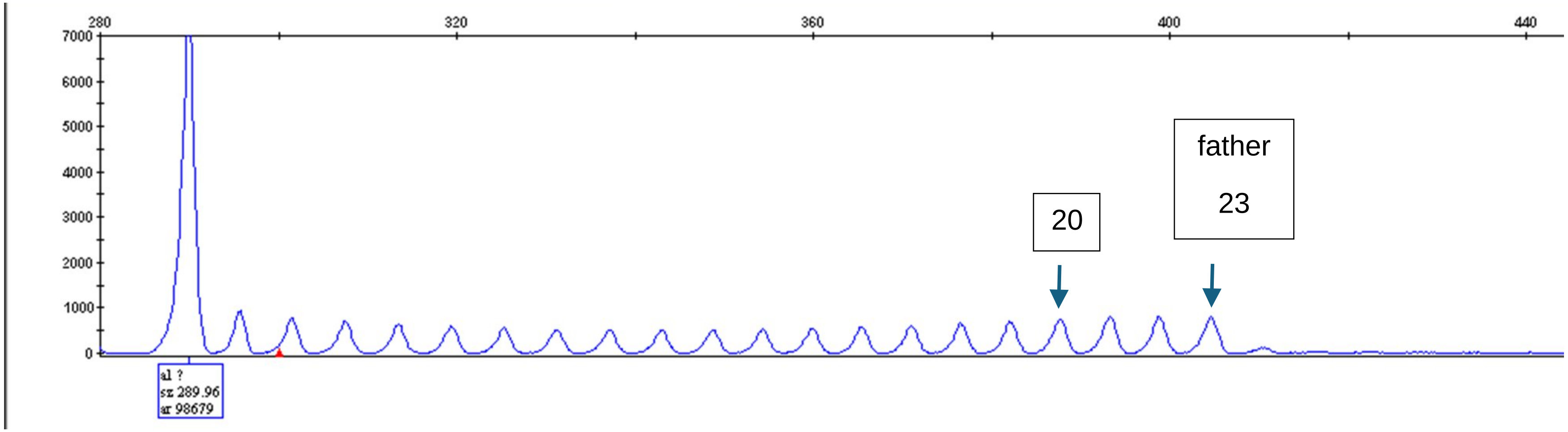
280
320
360
400
440
7000
6000
5000
4000
3000
2000
1000
0
al ?
sz 289.96
ar 98679
20
father
23

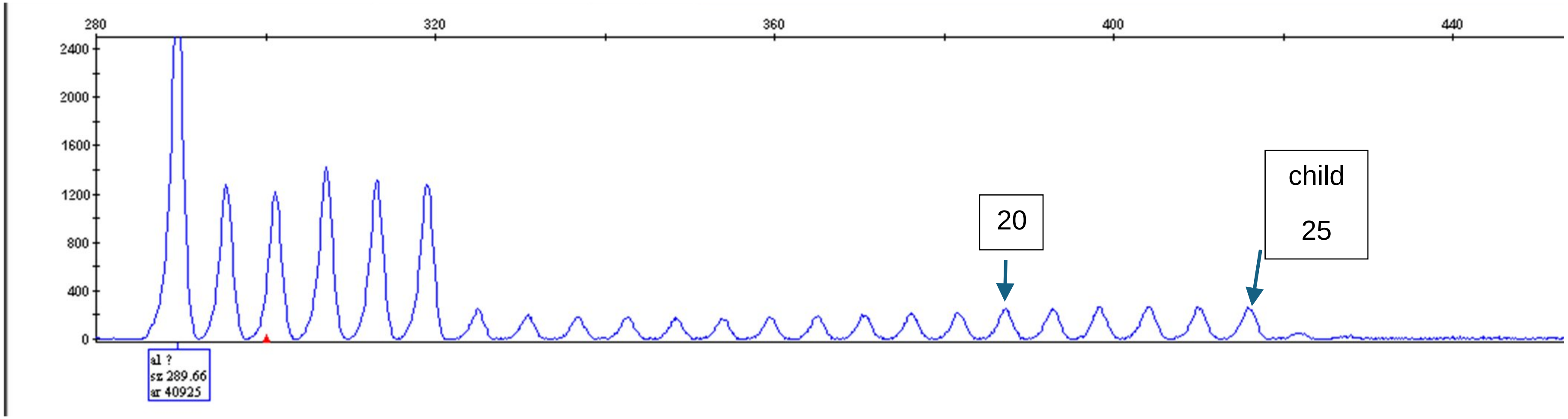
280
320
360
400
440
2400
2000
1600
1200
800
400
0
al ?
sz 289.66
ar 40925
20
child
25

**Supplemental Figure 2. AmplideX RP-PCR results for parent-offspring pairs (highest peak genotyping)**

fid: 12

**Parent**

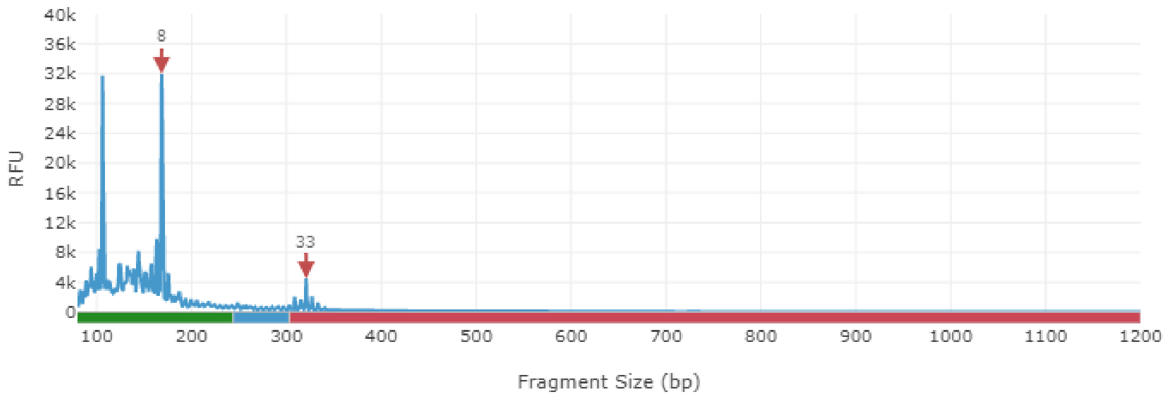


**Offspring**

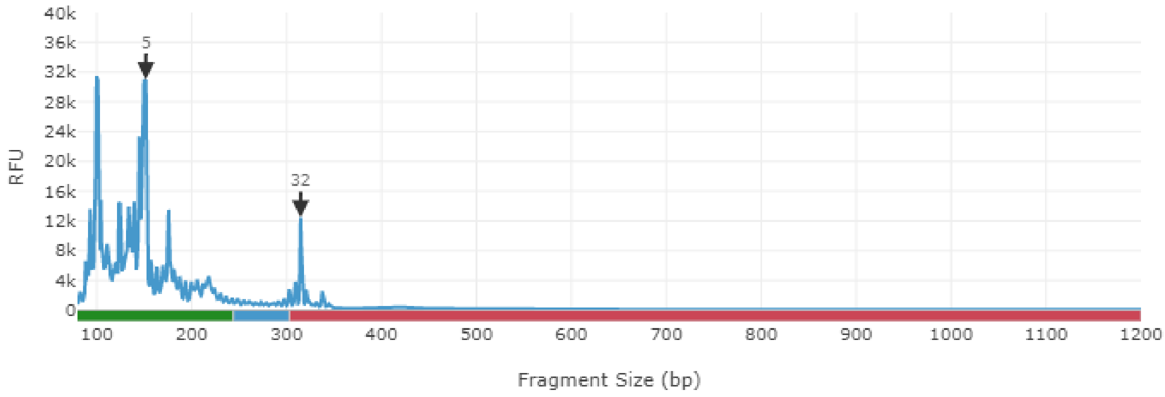

**Parent**

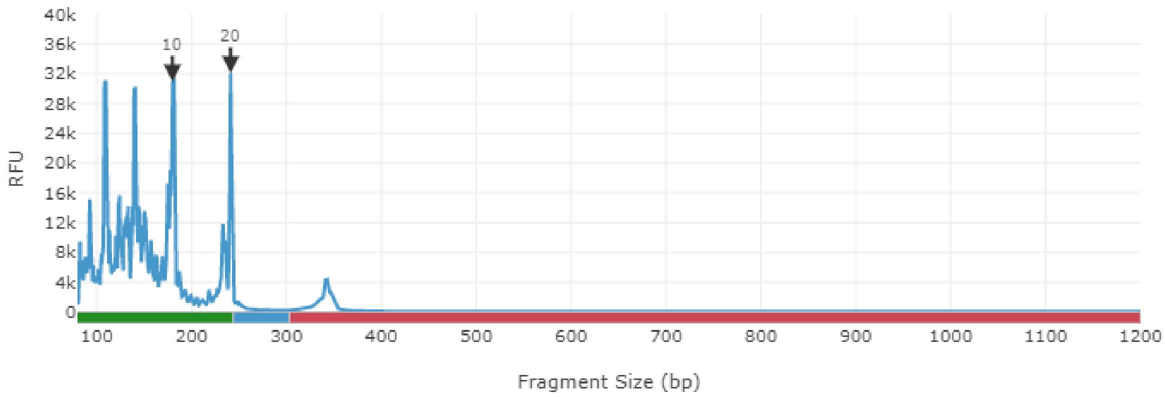


**Offspring**

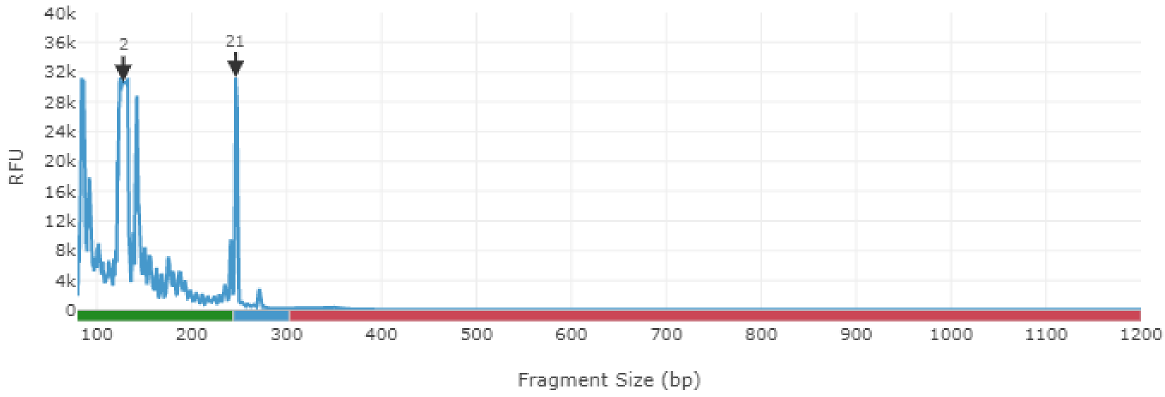

**Parent**

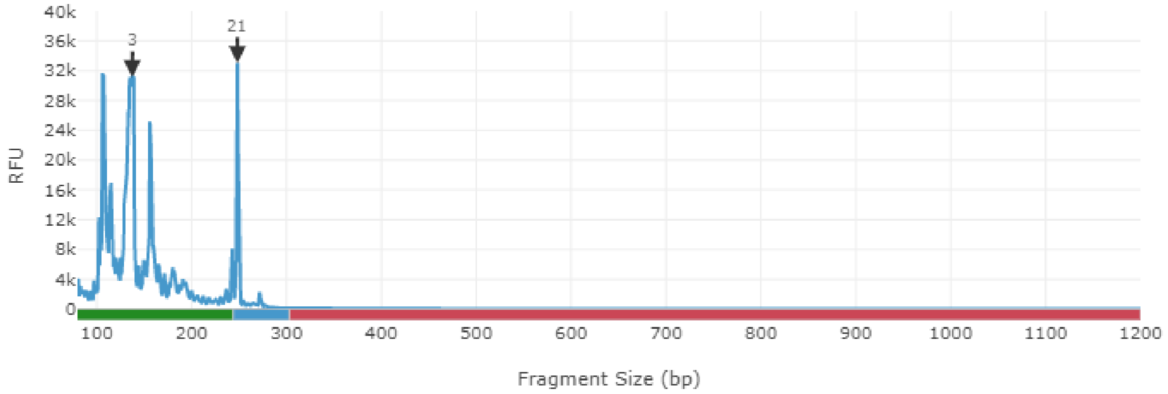


**Offspring**

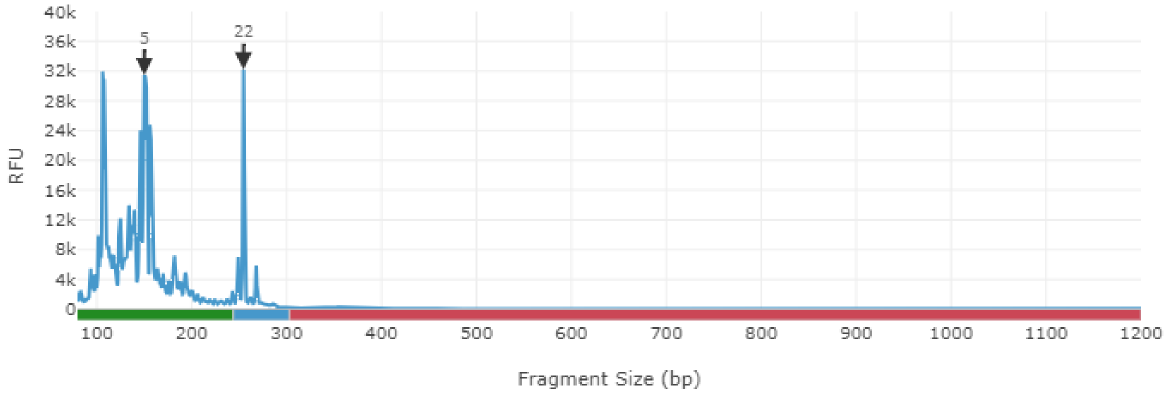

**Parent**

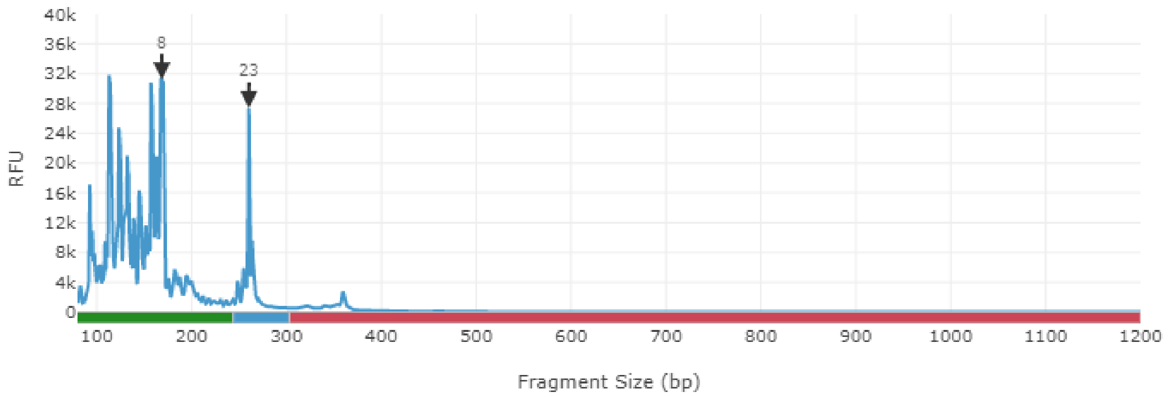


**Offspring**

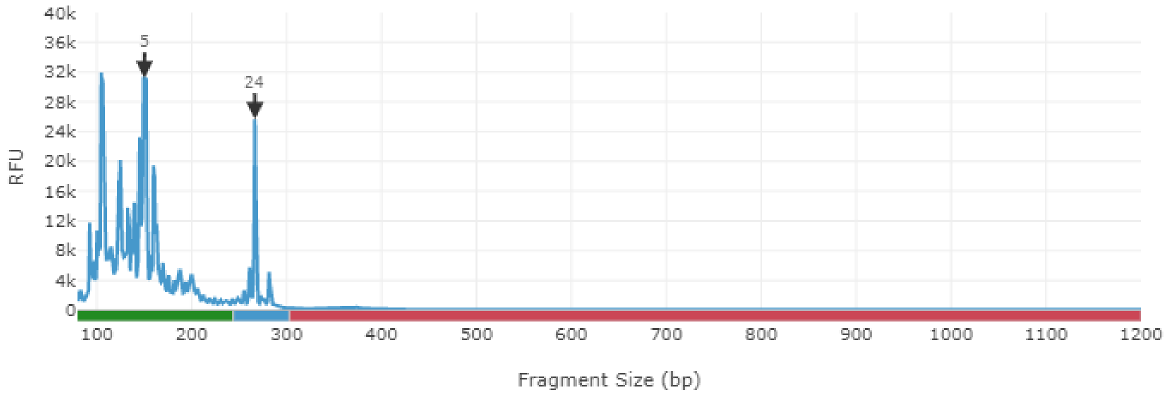

**Parent**

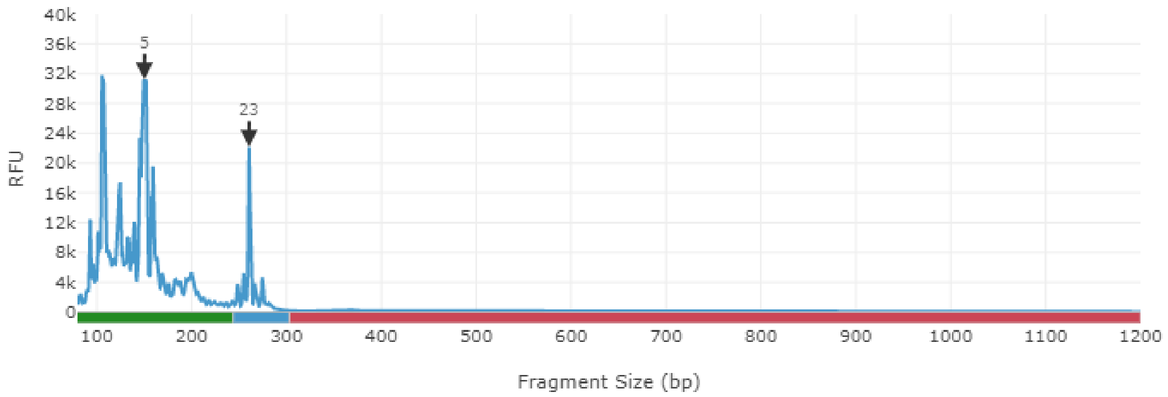


**Offspring**

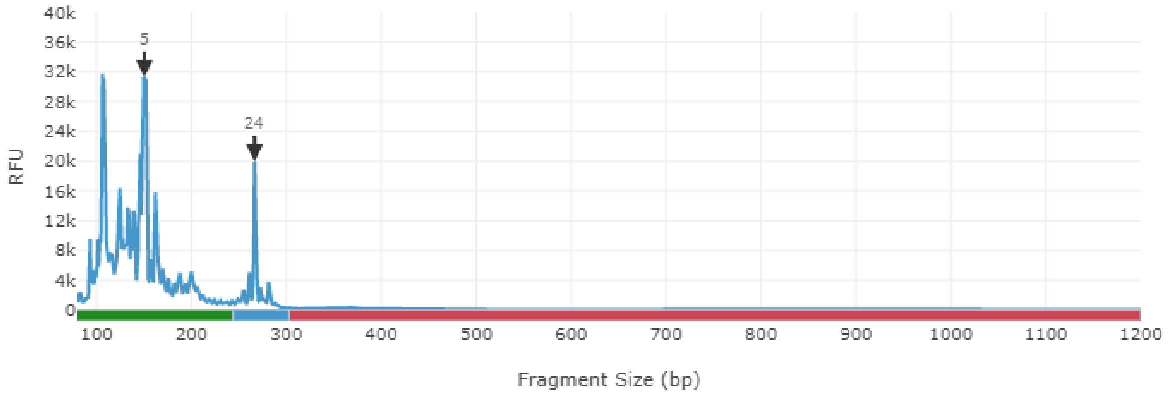

**Parent**

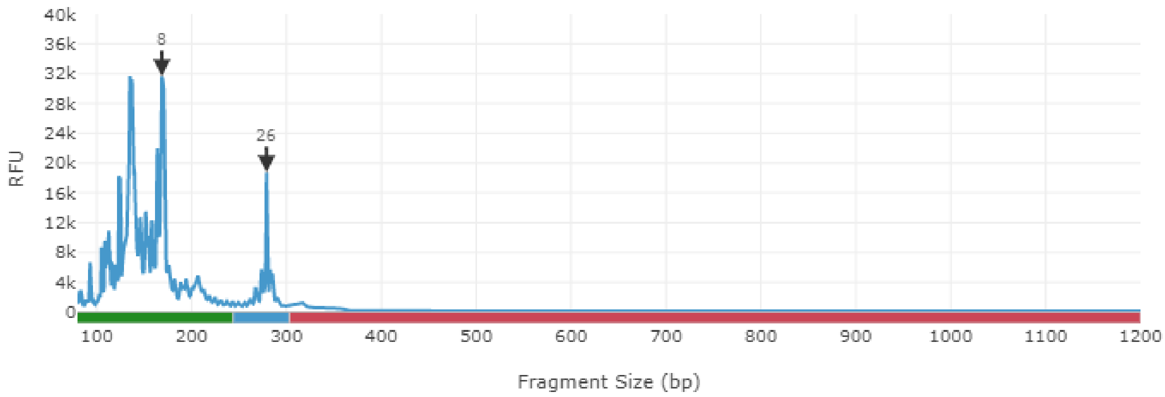


**Offspring**

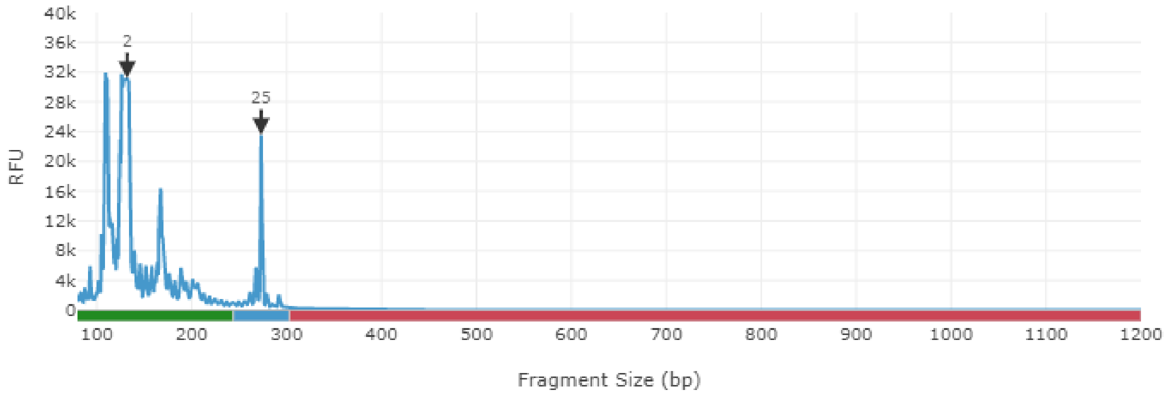

**Parent**

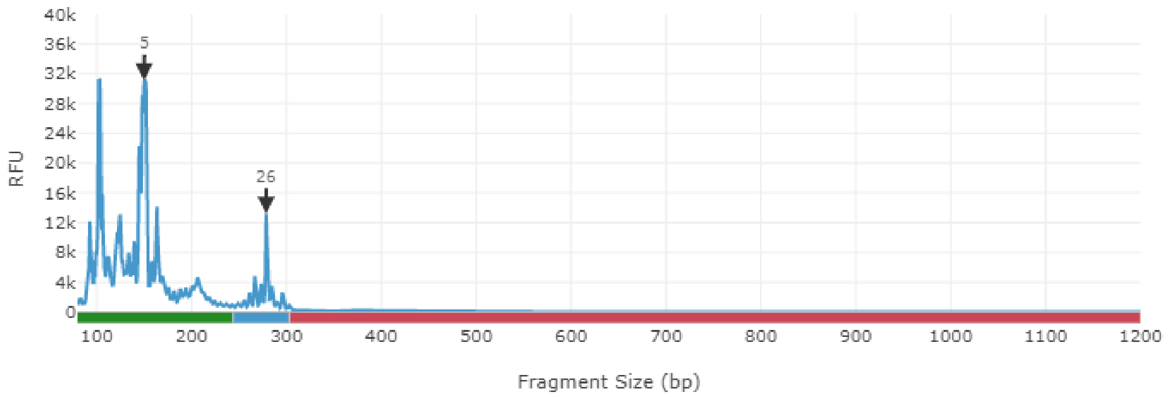


**Offspring**

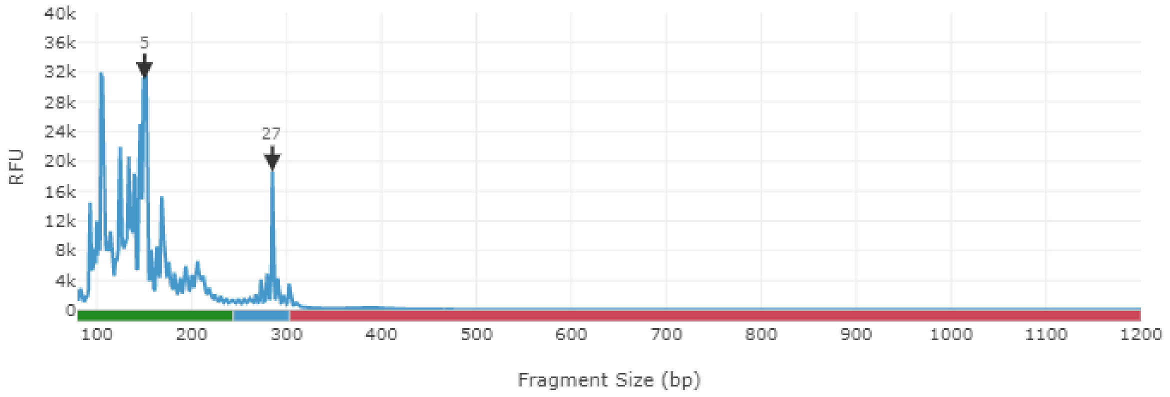

**Parent**

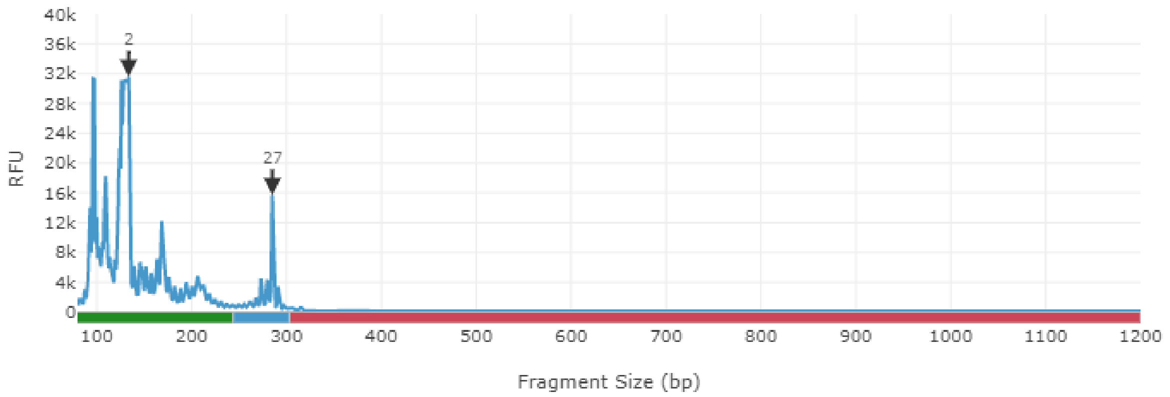


**Offspring**

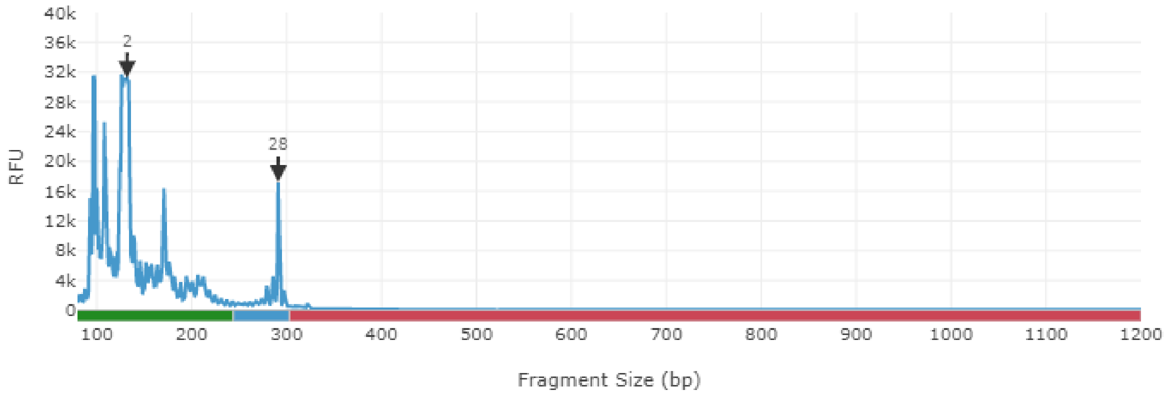

**Parent**

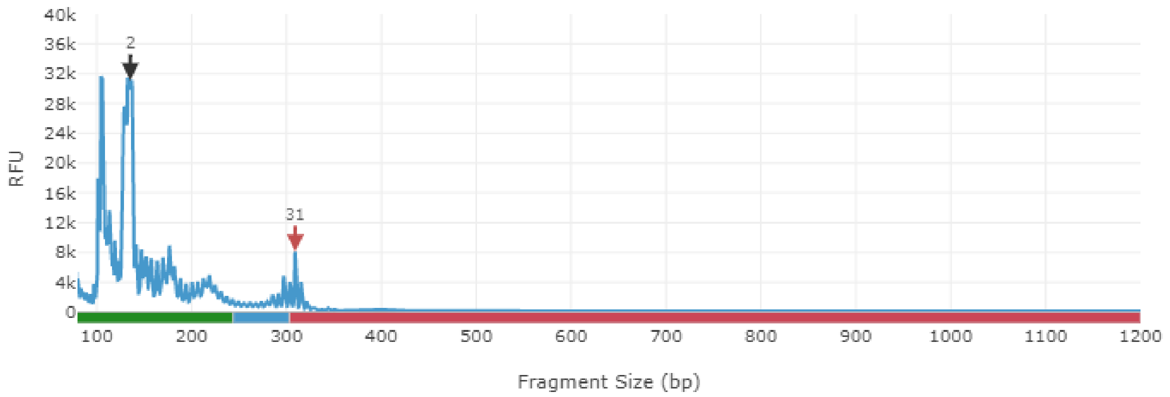


**Offspring**

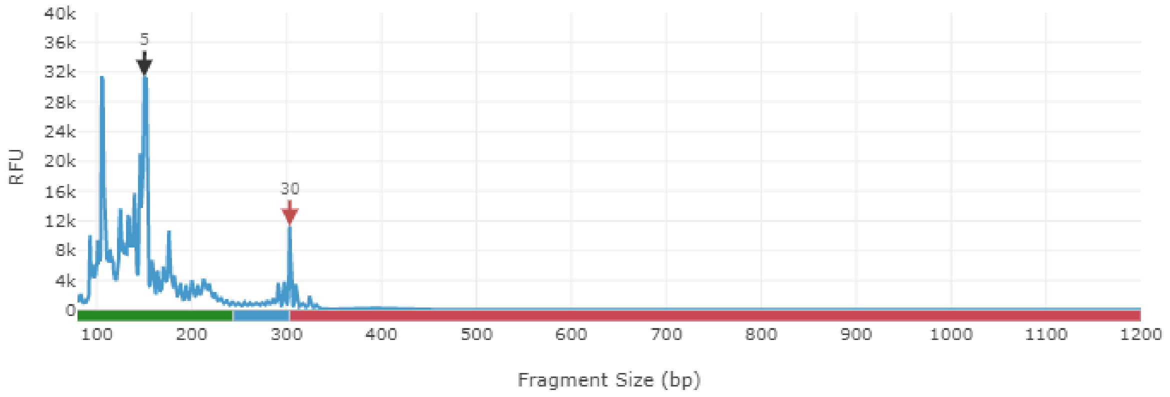

**Parent**

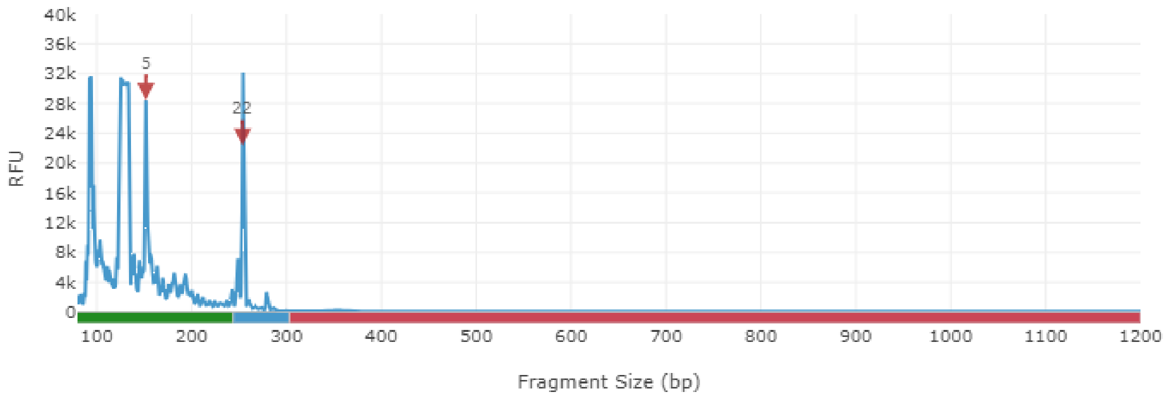


**Offspring**

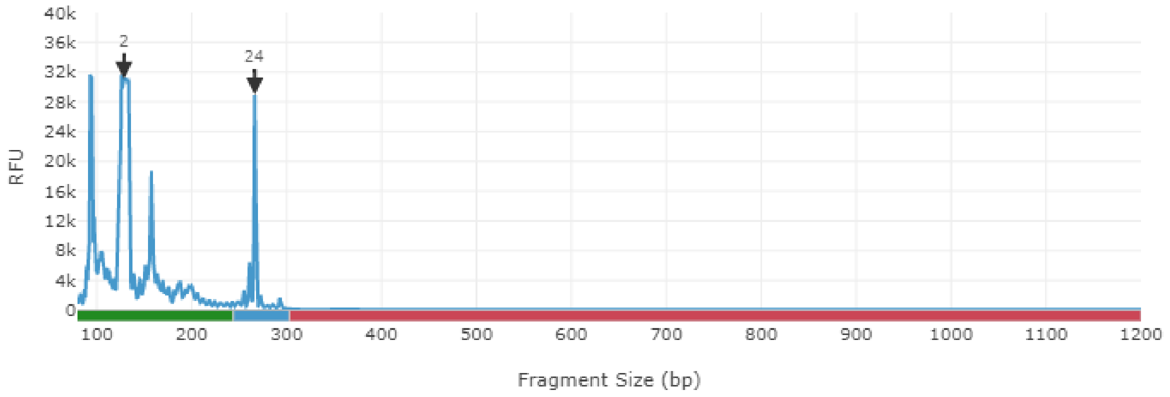

**Parent**

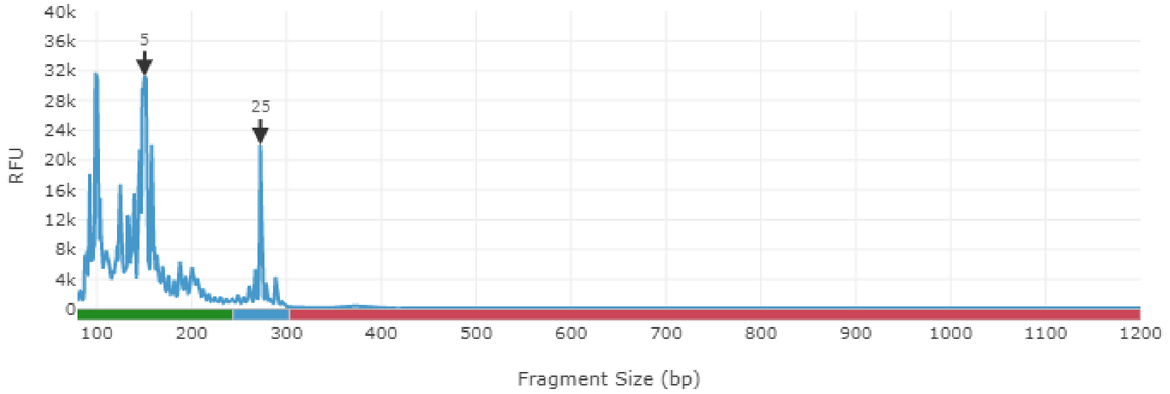


**Offspring**

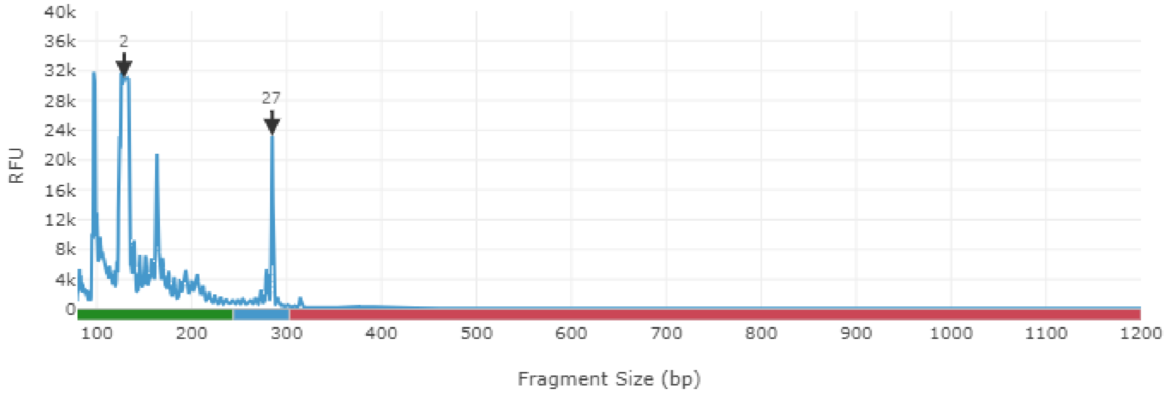

**Parent**

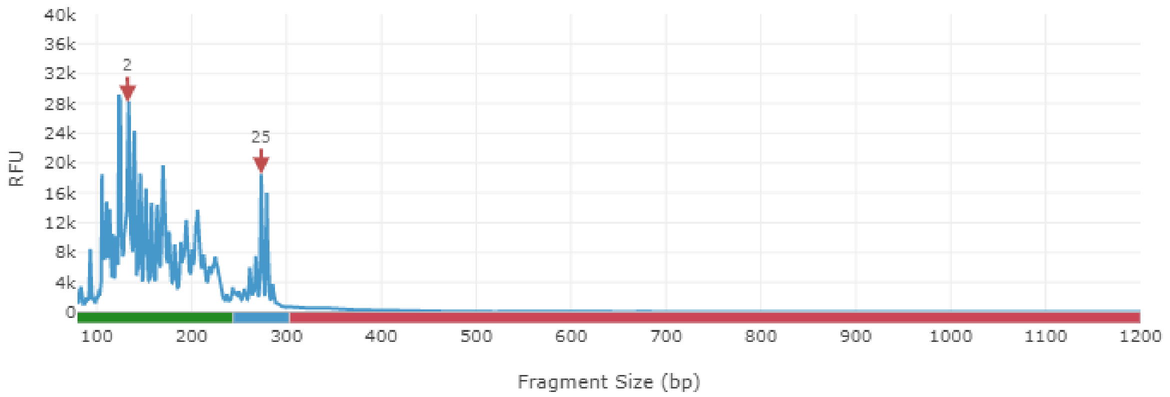


**Offspring**

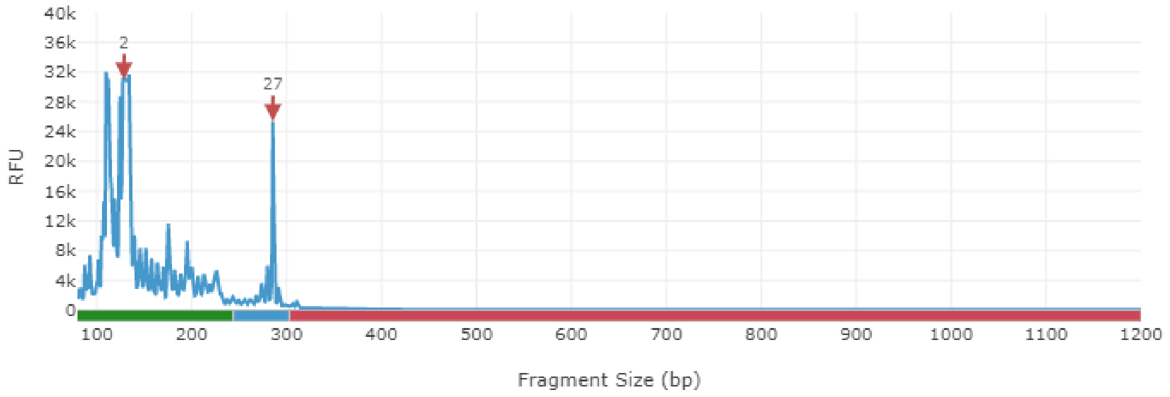

**Parent**

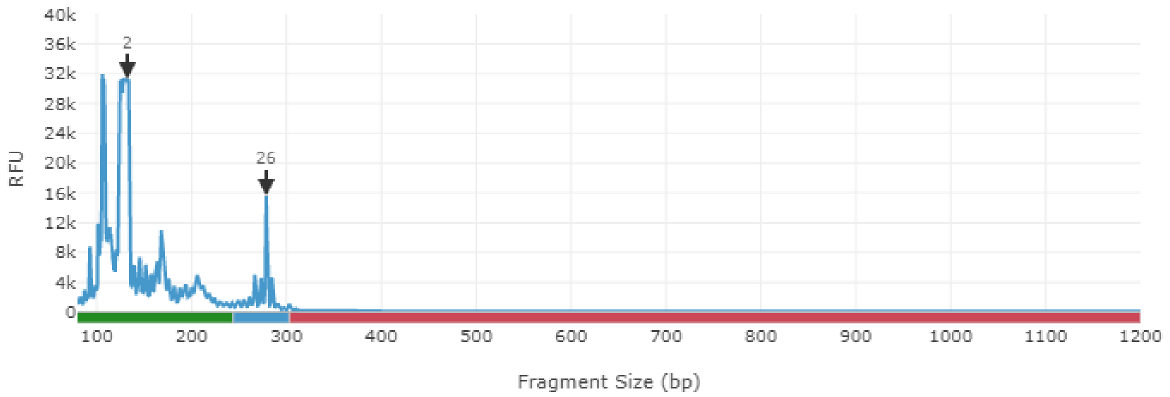


**Offspring**

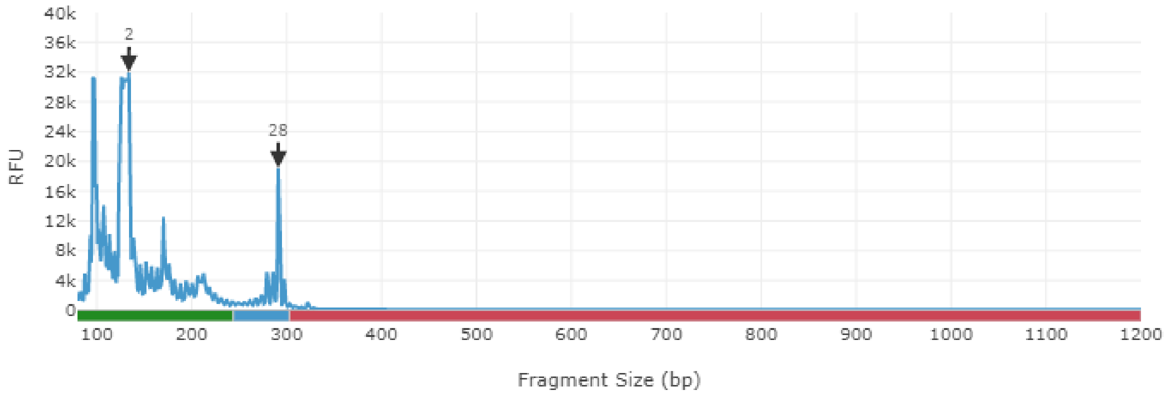

**Parent**

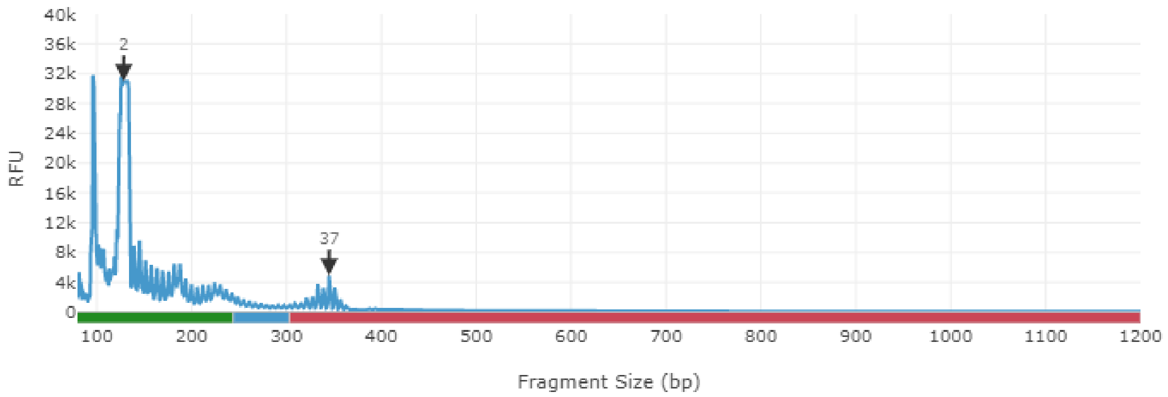


**Offspring**

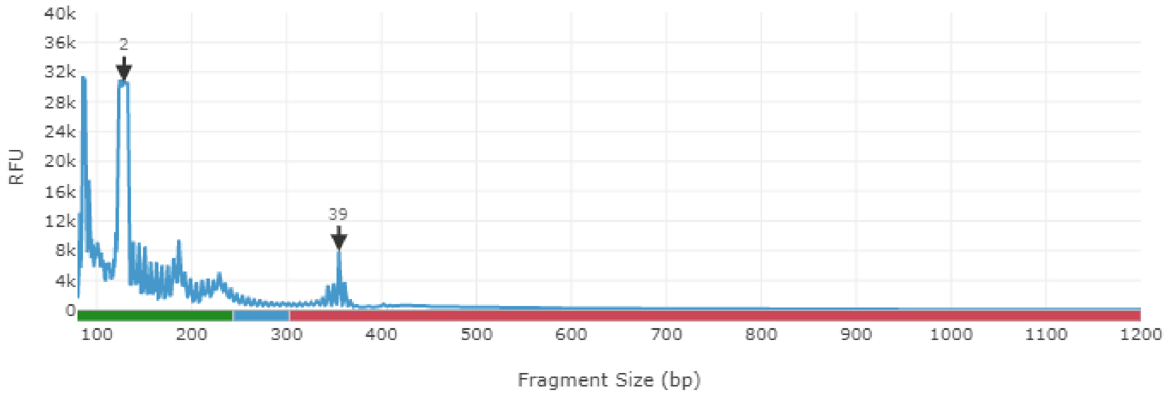

**Parent**

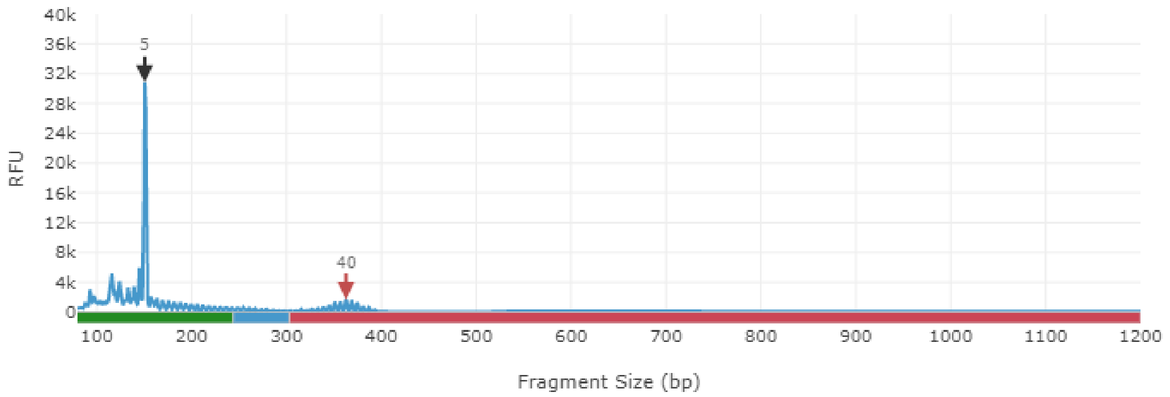


**Offspring**

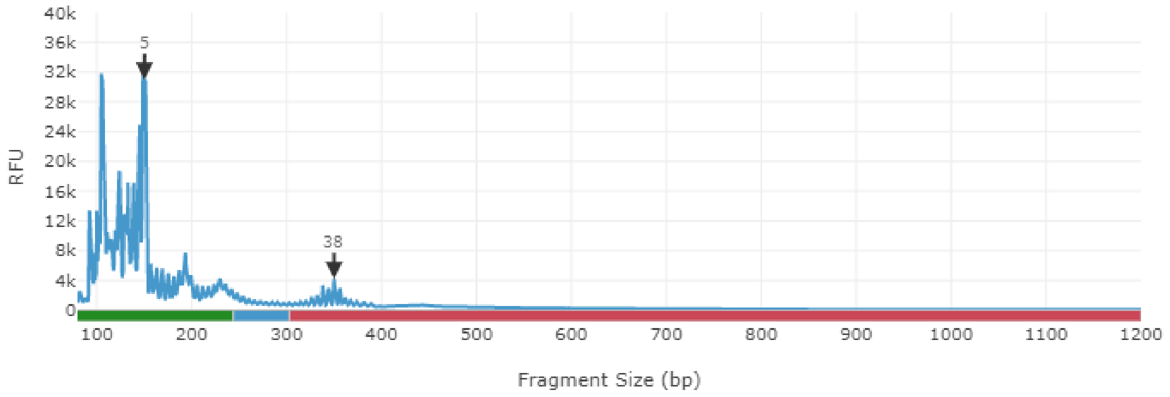

**Parent**

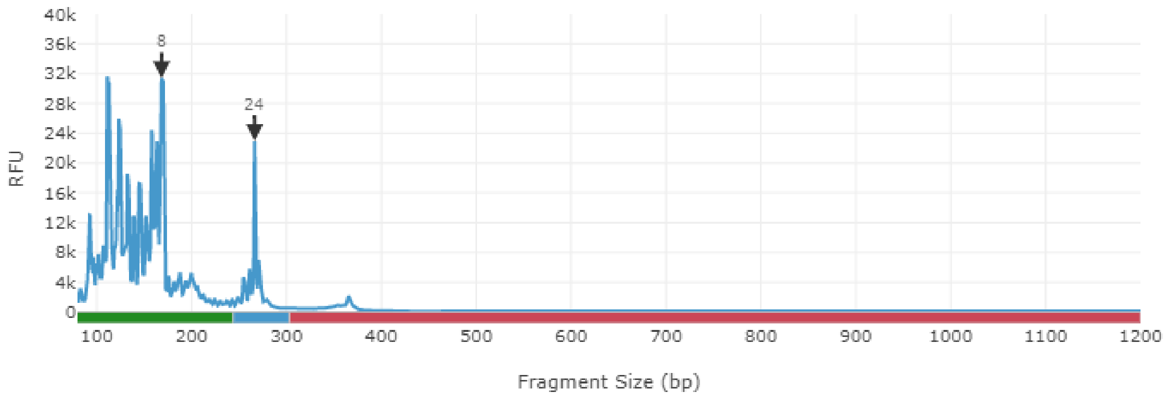


**Offspring**

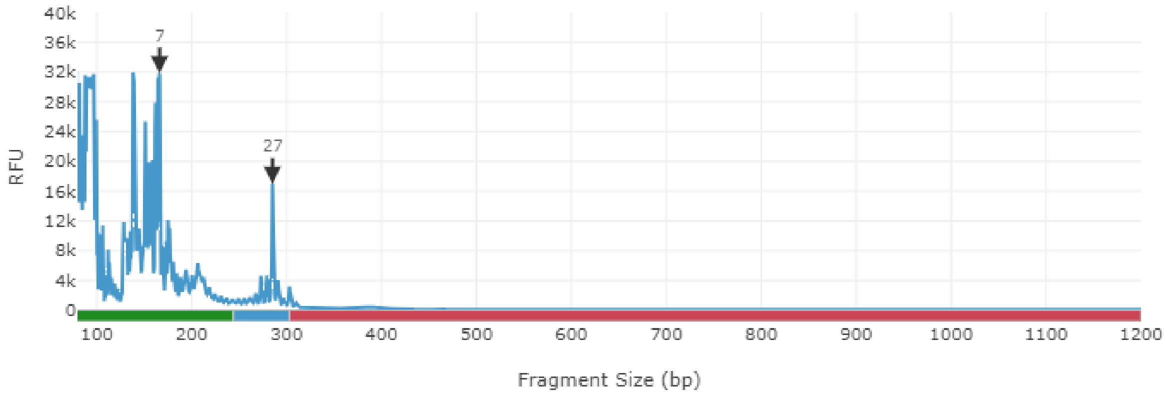

**Parent**

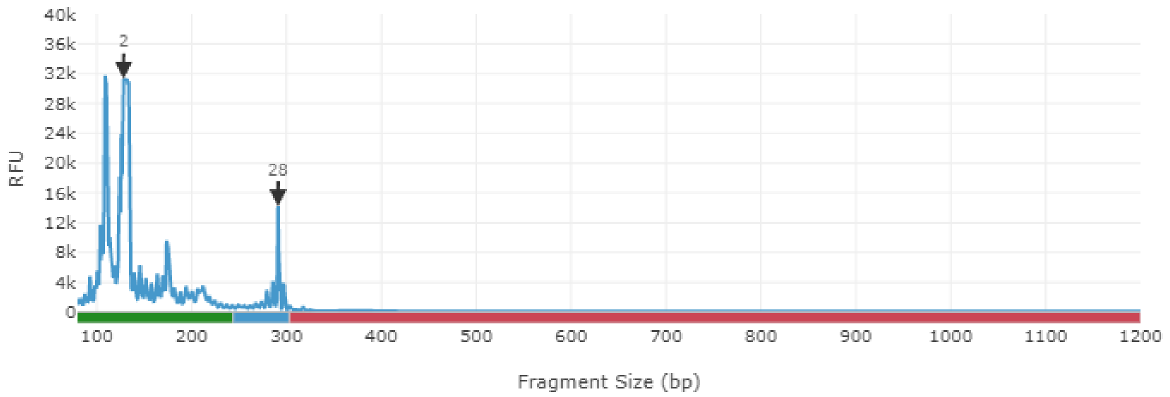


**Offspring**

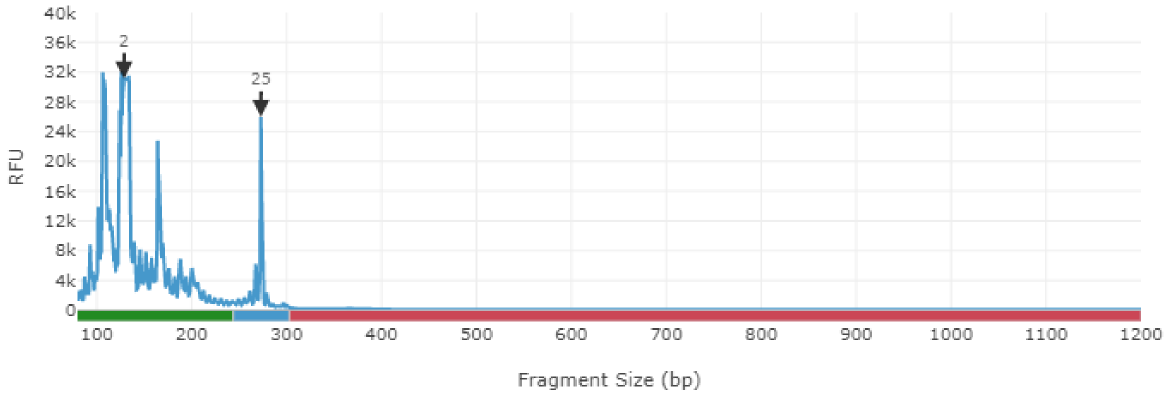

**Parent**

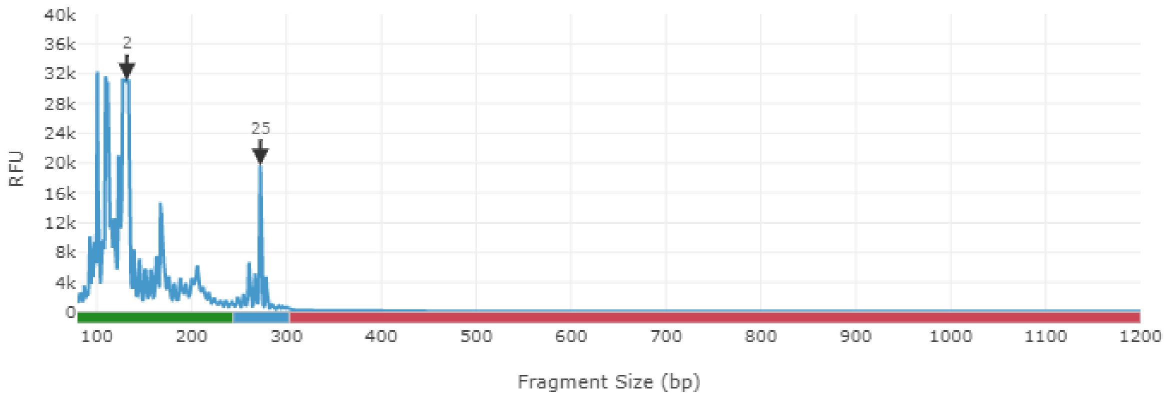


**Offspring**

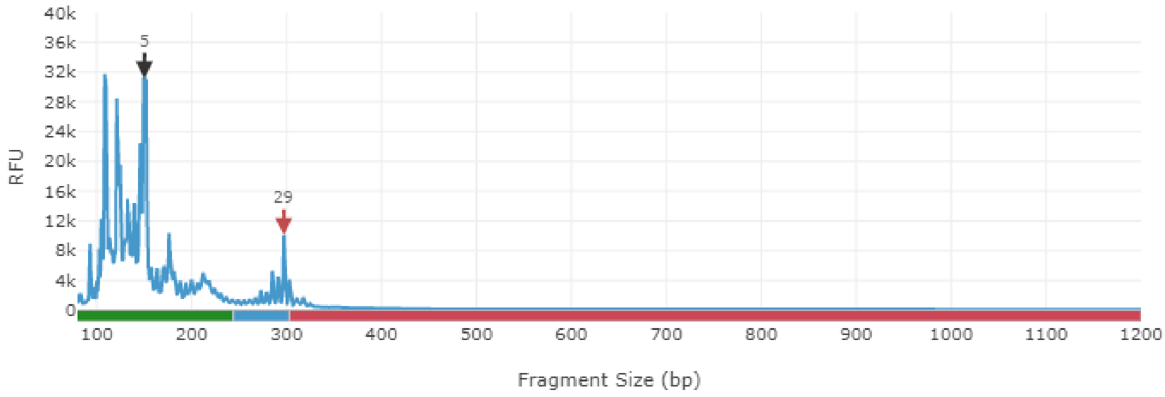

**Parent**

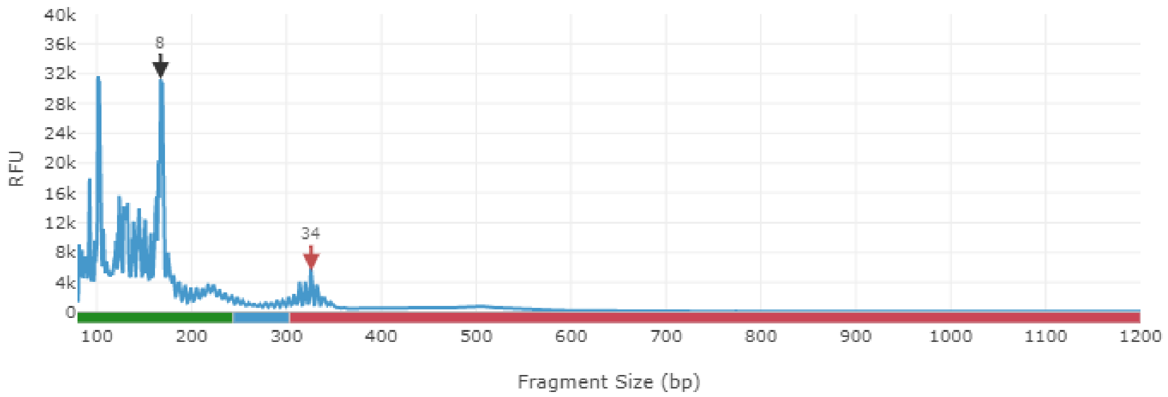


**Offspring**

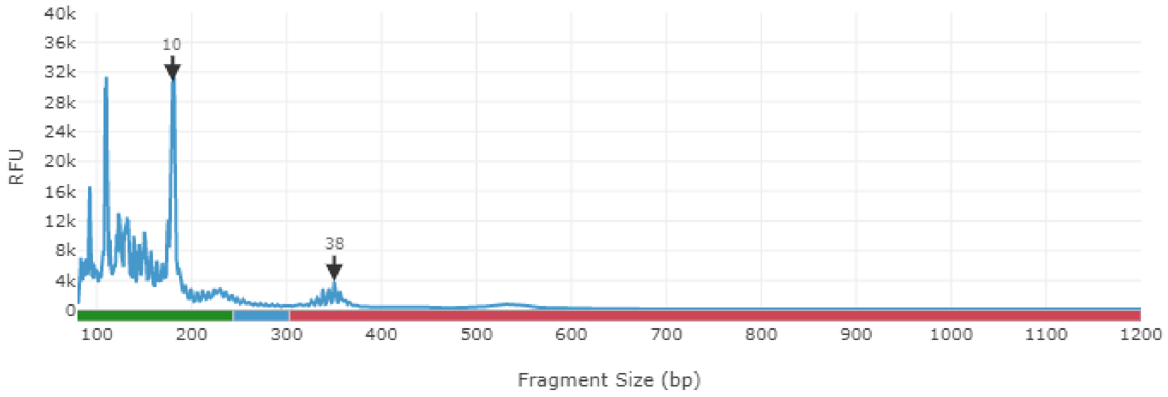

**Parent**

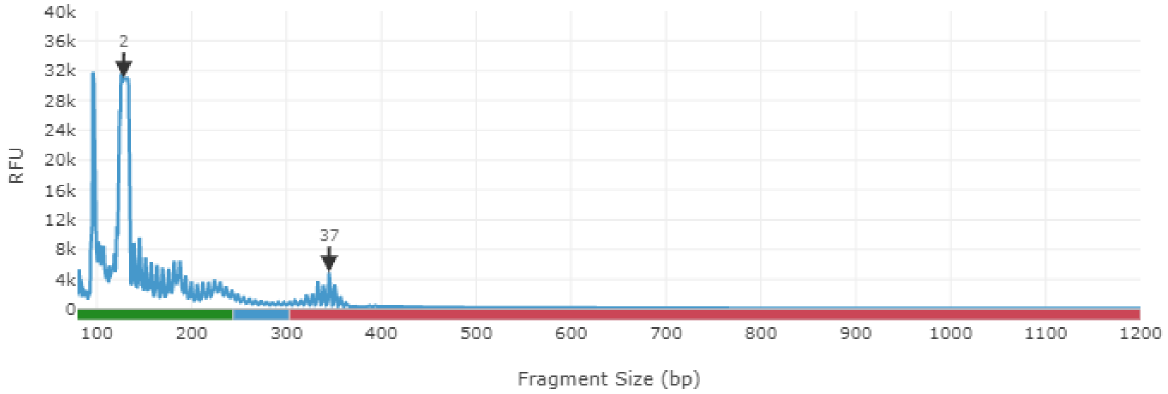


**Offspring**

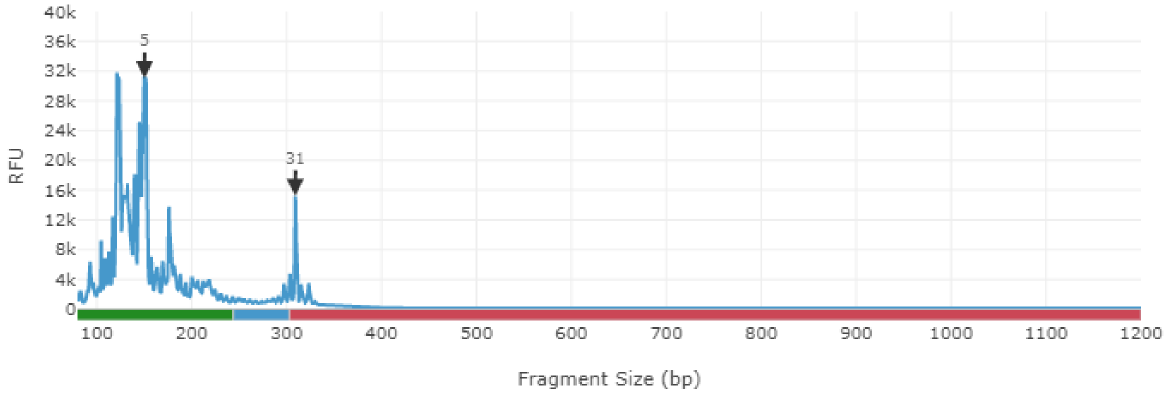

**Parent**

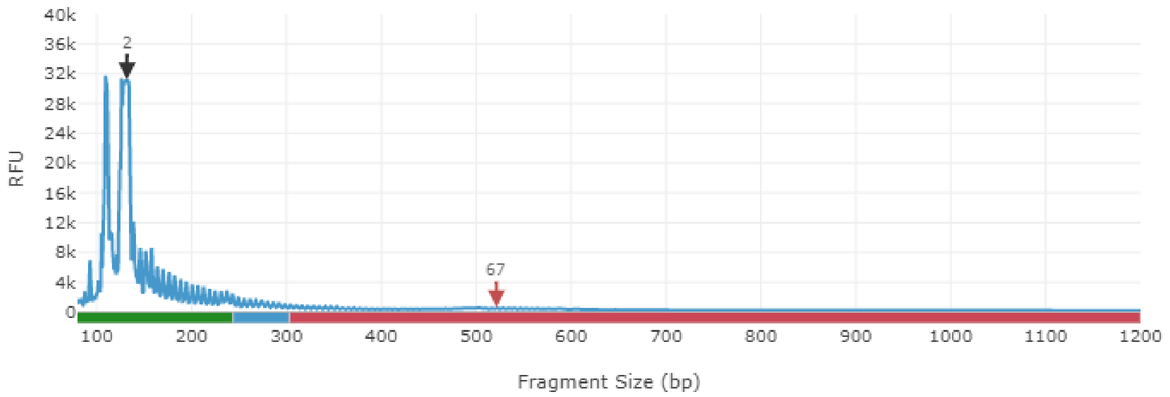


zoomed parent, highest peak

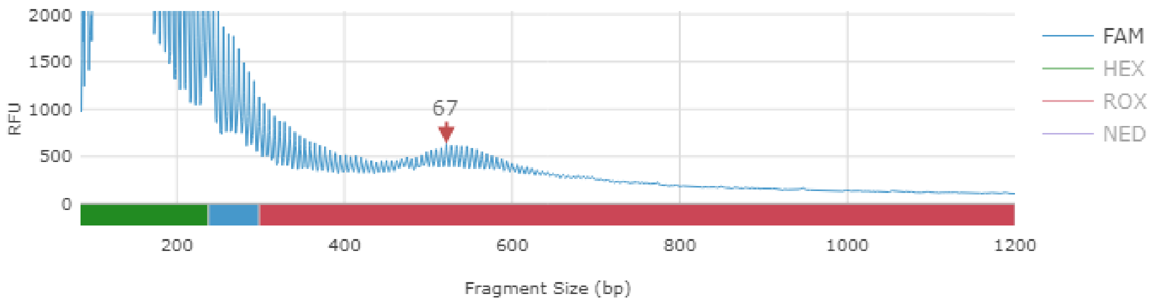


**Offspring**

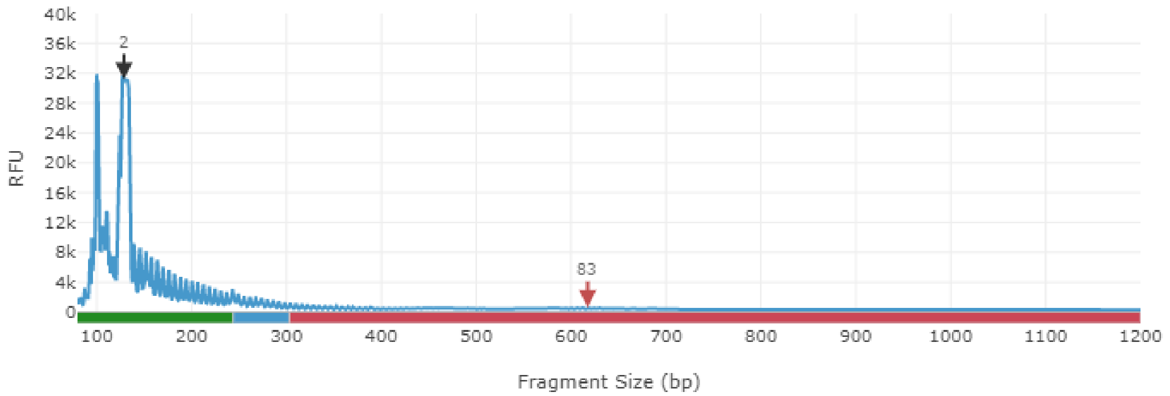


zoomed offspring, highest peak

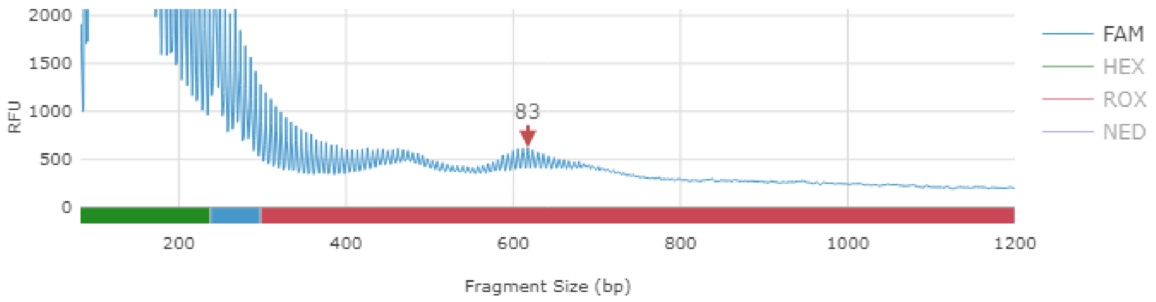

**Parent**

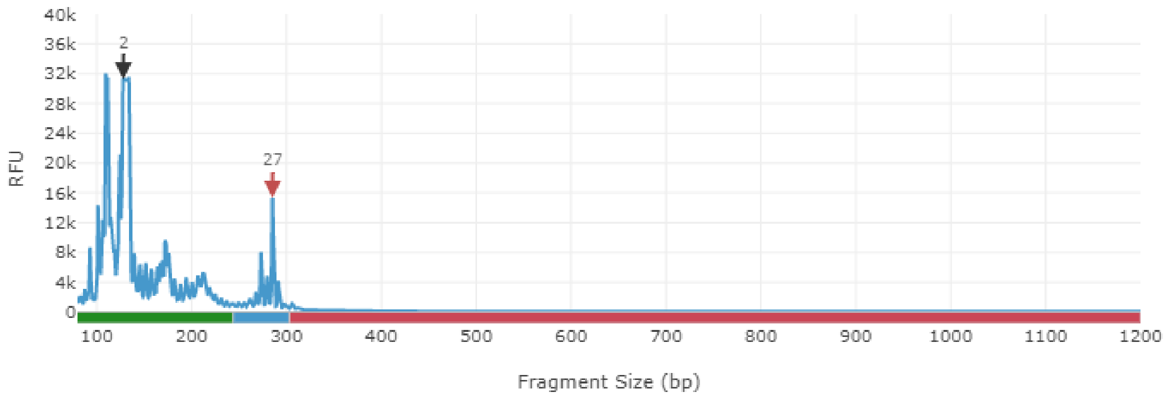


**Offspring**

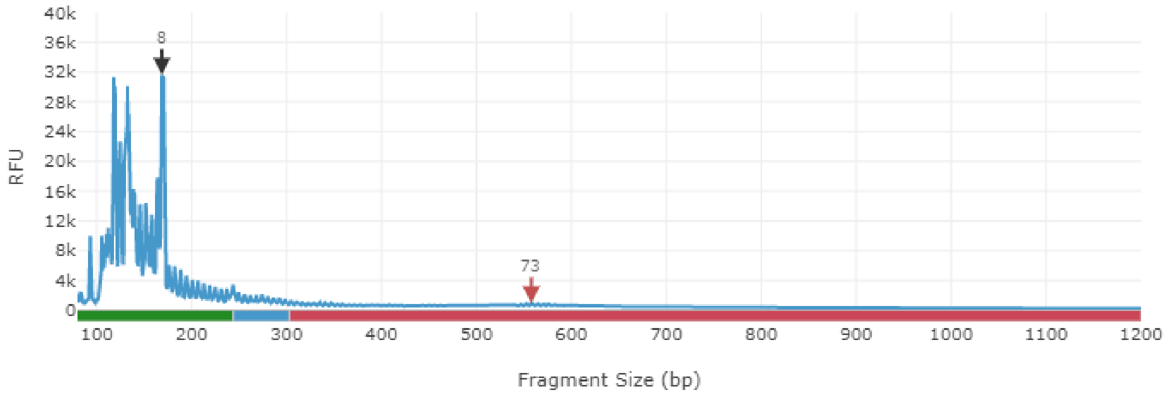


zoomed offspring, highest peak

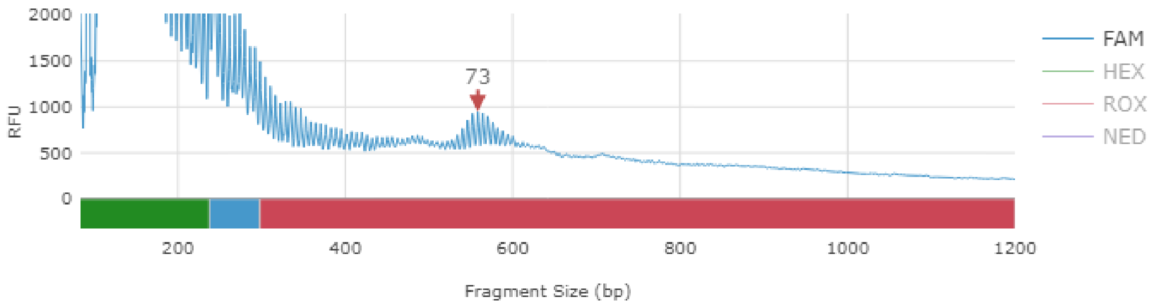

**Parent**

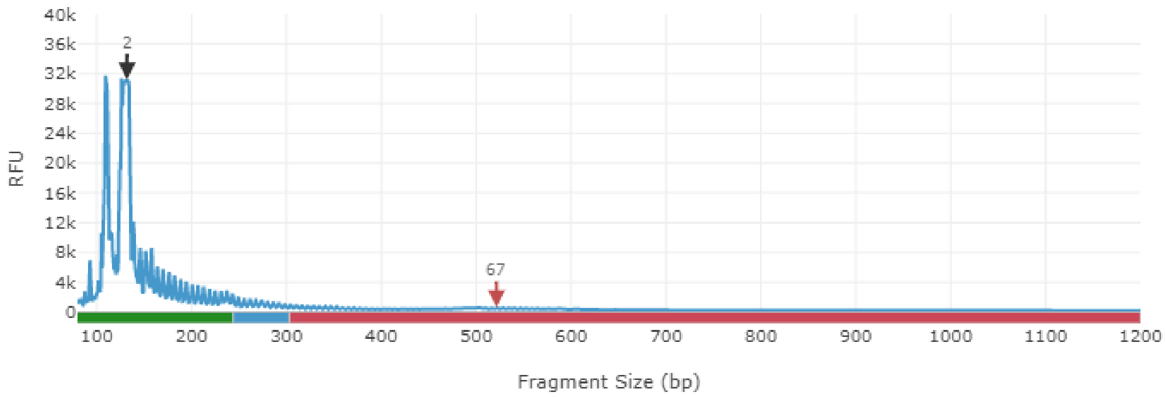


zoomed parent, highest peak

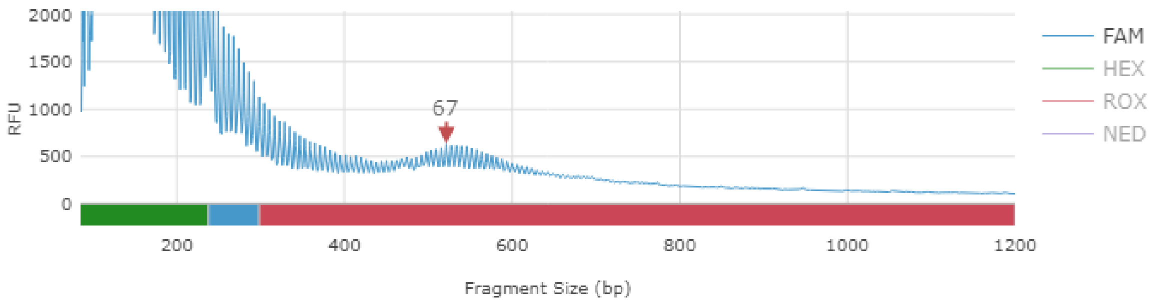


**Offspring**

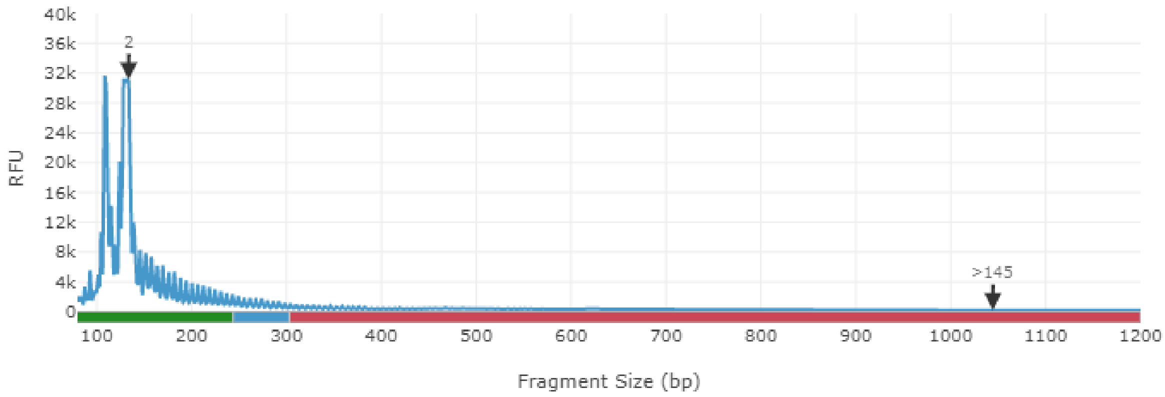


zoomed offspring, highest peak

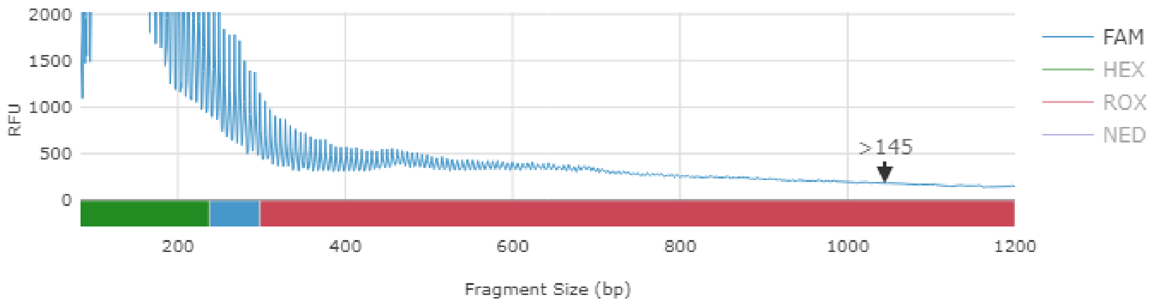
