## Supplemental Tables for "Intergenerational instability of the C9orf72 hexanucleotide repeat"

**Table 1. RP-PCR genotypes of the longer *C9orf72* GGGGCC allele**

The label `exp` denotes a repeat expansion. The suffix `in_house` denotes the "standard" lab RP-PCR while `amplidex` denotes the commercial and certified AmplideX RP-PCR kit.
The suffix `combined` denotes the consensus of `in-house` and `amplidex` genotypes where the standard RP-PCR genotypes were corrected to AmplideX genotypes if required.

| fid | parent_allele_in_house | child_allele_in_house | parent_allele_amplidex | child_allele_amplidex | change_in_house | change_amplidex | change_combined | parent_allele_combined |
|---|---|---|---|---|---|---|---|---|
| 3 | na | 14 | na | na | na | na | na | na |
| 3 | na | 14 | na | na | na | na | na | na |
| 17 | na | 30 | na | 31 | na | na | na | na |
| 32 | na | 11 | na | na | na | na | na | na |
| 33 | 11 | na | na | NA | na | na | na | na |
| 35 | 14 | na | na | 14 | na | na | na | na |
| 7 | 27 | na | 27 | na | na | na | na | na |
| 17 | 31 | na | 31 | na | na | na | na | na |
| 17 | 31 | na | 31 | na | na | na | na | na |
| 27 | 100 | na | 100 | na | na | na | na | na |
| 26 | na | na | na | na | 0 | na | 0 | na |
| 26 | na | na | na | na | 0 | na | 0 | na |
| 36 | 6 | 6 | na | na | 0 | na | 0 | 6 |
| 47 | 6 | 6 | na | na | 0 | na | 0 | 6 |
| 73 | 6 | 6 | na | na | 0 | na | 0 | 6 |
| 38 | 7 | 7 | na | na | 0 | na | 0 | 7 |
| 37 | 8 | 8 | na | na | 0 | na | 0 | 8 |
| 37 | 8 | 8 | na | na | 0 | na | 0 | 8 |
| 54 | 8 | 8 | na | na | 0 | na | 0 | 8 |
| 67 | 8 | 8 | na | na | 0 | na | 0 | 8 |
| 77 | 8 | 8 | na | na | 0 | na | 0 | 8 |
| 78 | 8 | 8 | na | na | 0 | na | 0 | 8 |
| 88 | 10 | 10 | na | na | 0 | na | 0 | 10 |
| 39 | 11 | 11 | na | na | 0 | na | 0 | 11 |
| 48 | 11 | 11 | na | na | 0 | na | 0 | 11 |
| 53 | 11 | 11 | na | na | 0 | na | 0 | 11 |
| 53 | 11 | 11 | na | na | 0 | na | 0 | 11 |
| 63 | 11 | 11 | na | na | 0 | na | 0 | 11 |
| 64 | 11 | 11 | na | na | 0 | na | 0 | 11 |
| 66 | 11 | 11 | na | na | 0 | na | 0 | 11 |
| 69 | 11 | 11 | na | na | 0 | na | 0 | 11 |
| 99 | 11 | 11 | na | na | 0 | na | 0 | 11 |
| 99 | 11 | 11 | na | na | 0 | na | 0 | 11 |
| 106 | 11 | 11 | na | na | 0 | na | 0 | 11 |

| | | | | | | | | |
|---|---|---|---|---|---|---|---|---|
| 113 | 11 | 11 | na | na | 0 | na | 0 | 11 |
| 123 | 11 | 11 | na | na | 0 | na | 0 | 11 |
| 40 | 13 | 13 | na | na | 0 | na | 0 | 13 |
| 40 | 13 | 13 | na | na | 0 | na | 0 | 13 |
| 49 | 13 | 13 | na | na | 0 | na | 0 | 13 |
| 49 | 13 | 13 | na | na | 0 | na | 0 | 13 |
| 49 | 13 | 13 | na | na | 0 | na | 0 | 13 |
| 80 | 13 | 13 | na | na | 0 | na | 0 | 13 |
| 94 | 13 | 13 | na | na | 0 | na | 0 | 13 |
| 127 | 13 | 13 | na | na | 0 | na | 0 | 13 |
| 43 | 14 | 14 | na | na | 0 | na | 0 | 14 |
| 43 | 14 | 14 | na | na | 0 | na | 0 | 14 |
| 61 | 14 | 14 | na | na | 0 | na | 0 | 14 |
| 61 | 14 | 14 | na | na | 0 | na | 0 | 14 |
| 65 | 14 | 14 | na | na | 0 | na | 0 | 14 |
| 75 | 14 | 14 | na | na | 0 | na | 0 | 14 |
| 75 | 14 | 14 | na | na | 0 | na | 0 | 14 |
| 75 | 14 | 14 | na | na | 0 | na | 0 | 14 |
| 82 | 14 | 14 | na | na | 0 | na | 0 | 14 |
| 86 | 14 | 14 | na | na | 0 | na | 0 | 14 |
| 89 | 14 | 14 | na | na | 0 | na | 0 | 14 |
| 96 | 14 | 14 | na | na | 0 | na | 0 | 14 |
| 119 | 14 | 14 | na | na | 0 | na | 0 | 14 |
| 121 | 14 | 14 | na | na | 0 | na | 0 | 14 |
| 124 | 14 | 14 | na | na | 0 | na | 0 | 14 |
| 120 | 15 | 15 | na | na | 0 | na | 0 | 15 |
| 102 | 16 | 16 | na | na | 0 | na | 0 | 16 |
| 102 | 16 | 16 | na | na | 0 | na | 0 | 16 |
| 102 | 16 | 16 | na | na | 0 | na | 0 | 16 |
| 103 | 16 | 16 | na | na | 0 | na | 0 | 16 |
| 103 | 16 | 16 | na | na | 0 | na | 0 | 16 |
| 45 | 18 | 18 | na | na | 0 | na | 0 | 18 |
| 56 | 18 | 18 | na | na | 0 | na | 0 | 18 |
| 56 | 18 | 18 | na | na | 0 | na | 0 | 18 |
| 59 | 18 | 18 | na | na | 0 | na | 0 | 18 |
| 59 | 18 | 18 | na | na | 0 | na | 0 | 18 |
| 70 | 18 | 18 | na | na | 0 | na | 0 | 18 |
| 87 | 18 | 18 | na | na | 0 | na | 0 | 18 |
| 91 | 18 | 18 | na | na | 0 | na | 0 | 18 |

| | | | | | | | | |
|---|---|---|---|---|---|---|---|---|
| 114 | 18 | 18 | na | na | 0 | na | 0 | 18 |
| 114 | 18 | 18 | na | na | 0 | na | 0 | 18 |
| 85 | 19 | 19 | na | na | 0 | na | 0 | 19 |
| 11 | 20 | 20 | na | na | 0 | na | 0 | 20 |
| 11 | 20 | 20 | na | na | 0 | na | 0 | 20 |
| 51 | 20 | 20 | na | na | 0 | na | 0 | 20 |
| 55 | 20 | 20 | na | na | 0 | na | 0 | 20 |
| 60 | 20 | 20 | na | na | 0 | na | 0 | 20 |
| 98 | 20 | 20 | na | na | 0 | na | 0 | 20 |
| 112 | 20 | 20 | na | na | 0 | na | 0 | 20 |
| 112 | 20 | 20 | na | na | 0 | na | 0 | 20 |
| 52 | 21 | 21 | na | na | 0 | na | 0 | 21 |
| 68 | 21 | 21 | na | na | 0 | na | 0 | 21 |
| 84 | 21 | 21 | na | na | 0 | na | 0 | 21 |
| 101 | 21 | 21 | na | na | 0 | na | 0 | 21 |
| 110 | 21 | 21 | na | na | 0 | na | 0 | 21 |
| 111 | 21 | 21 | na | na | 0 | na | 0 | 21 |
| 115 | 21 | 21 | na | na | 0 | na | 0 | 21 |
| 115 | 21 | 21 | na | na | 0 | na | 0 | 21 |
| 19 | 21 | 21 | 21 | na | 0 | na | 0 | 21 |
| 19 | 21 | 5 | 21 | 21 | -16 | 0 | 0 | 21 |
| 22 | 21 | 21 | 21 | na | 0 | na | 0 | 21 |
| 22 | 21 | 21 | 21 | na | 0 | na | 0 | 21 |
| 41 | 22 | 22 | na | na | 0 | na | 0 | 22 |
| 57 | 22 | 22 | na | na | 0 | na | 0 | 22 |
| 71 | 22 | 22 | na | na | 0 | na | 0 | 22 |
| 97 | 22 | 22 | na | na | 0 | na | 0 | 22 |
| 108 | 22 | 22 | na | na | 0 | na | 0 | 22 |
| 108 | 22 | 22 | na | na | 0 | na | 0 | 22 |
| 34 | 23 | 23 | na | na | 0 | na | 0 | 23 |
| 50 | 23 | 23 | na | na | 0 | na | 0 | 23 |
| 79 | 23 | 23 | na | na | 0 | na | 0 | 23 |
| 117 | 23 | 23 | na | na | 0 | na | 0 | 23 |
| 117 | 23 | 23 | na | na | 0 | na | 0 | 23 |
| 118 | 23 | 23 | na | na | 0 | na | 0 | 23 |
| 58 | 24 | 24 | na | na | 0 | na | 0 | 24 |
| 104 | 24 | 24 | na | na | 0 | na | 0 | 24 |
| 44 | 25 | 25 | na | na | 0 | na | 0 | 25 |
| 62 | 25 | 25 | na | na | 0 | na | 0 | 25 |

| | | | | | | | | |
|---|---|---|---|---|---|---|---|---|
| 126 | 25 | 25 | na | na | 0 | na | 0 | 25 |
| 128 | 25 | 25 | na | na | 0 | na | 0 | 25 |
| 23 | 26 | 26 | 26 | na | 0 | na | 0 | 26 |
| 31 | 26 | na | 26 | 26 | na | 0 | 0 | 26 |
| 18 | 27 | 27 | 27 | na | 0 | na | 0 | 27 |
| 18 | 27 | 27 | 27 | na | 0 | na | 0 | 27 |
| 7 | 27 | 26 | 27 | 27 | -1 | 0 | 0 | 27 |
| 7 | 27 | 27 | 27 | na | 0 | na | 0 | 27 |
| 116 | 28 | 28 | na | na | 0 | na | 0 | 28 |
| 10 | 28 | 28 | 28 | na | 0 | na | 0 | 28 |
| 42 | 29 | 29 | na | na | 0 | na | 0 | 29 |
| 90 | 29 | 29 | na | na | 0 | na | 0 | 29 |
| 122 | 33 | 33 | na | na | 0 | na | 0 | 33 |
| 25 | 100 | 100 | 37 | 37 | 0 | 0 | 0 | 37 |
| 28 | na | 100 | 100 | 100 | na | 0 | 0 | exp |
| 46 | 100 | 100 | 100 | 100 | 0 | 0 | 0 | exp |
| 46 | 100 | 100 | 100 | 100 | 0 | 0 | 0 | exp |
| 74 | 100 | 100 | 100 | 100 | 0 | 0 | 0 | exp |
| 76 | 100 | 100 | 100 | 100 | 0 | 0 | 0 | exp |
| 81 | 100 | 100 | 100 | 100 | 0 | 0 | 0 | exp |
| 81 | 100 | 100 | 100 | 100 | 0 | 0 | 0 | exp |
| 81 | 100 | 100 | 100 | 100 | 0 | 0 | 0 | exp |
| 93 | 100 | 100 | 100 | 100 | 0 | 0 | 0 | exp |
| 100 | 100 | 100 | 100 | 100 | 0 | 0 | 0 | exp |
| 105 | 100 | 100 | 100 | 100 | 0 | 0 | 0 | exp |
| 107 | 100 | 100 | 100 | 100 | 0 | 0 | 0 | exp |
| 125 | 100 | 100 | 100 | 100 | 0 | 0 | 0 | exp |
| 12 | 6 | 32 | 33 | 32 | 24 | -1 | -1 | 32 |
| 14 | 20 | 21 | 20 | 21 | 1 | 1 | 1 | 20 |
| 22 | 21 | 22 | 21 | 22 | 1 | 1 | 1 | 21 |
| 8 | 23 | 22 | na | 22 | -1 | na | -1 | 23 |
| 24 | 23 | 24 | na | 24 | 1 | na | 1 | 23 |
| 1 | 23 | 24 | 23 | 24 | 1 | 1 | 1 | 23 |
| 6 | 23 | 24 | 23 | 24 | 1 | 1 | 1 | 23 |
| 9 | 26 | 25 | 26 | 25 | -1 | -1 | -1 | 26 |
| 15 | 26 | 27 | 26 | 27 | 1 | 1 | 1 | 26 |
| 18 | 27 | 28 | 27 | 28 | 1 | 1 | 1 | 27 |
| 4 | 31 | 30 | 31 | 30 | -1 | -1 | -1 | 31 |
| 21 | 22 | 24 | 22 | 24 | 2 | 2 | 2 | 22 |

| | | | | | | | | |
|---|---|---|---|---|---|---|---|---|
| 2 | 25 | 27 | 25 | 27 | 2 | 2 | 2 | 25 |
| 5 | 25 | 26 | 25 | 27 | 1 | 2 | 2 | 25 |
| 23 | 26 | 28 | 26 | 28 | 2 | 2 | 2 | 26 |
| 25 | 100 | 100 | 37 | 39 | 0 | 2 | 2 | 37 |
| 92 | 100 | 100 | 40 | 38 | 0 | -2 | -2 | 40 |
| 16 | 24 | 27 | 24 | 27 | 3 | 3 | 3 | 24 |
| 10 | 28 | 25 | 28 | 25 | -3 | -3 | -3 | 28 |
| 13 | 25 | 29 | 25 | 29 | 4 | 4 | 4 | 25 |
| 20 | 32 | 39 | 34 | 38 | 7 | 4 | 4 | 34 |
| 25 | 100 | 31 | 37 | 31 | -70 | -6 | -6 | 37 |
| 109 | 100 | 100 | 67 | 83 | 0 | 16 | 16 | 67 |
| 29 | na | 100 | 27 | 73 | na | 46 | 46 | 27 |
| 109 | 100 | 100 | 67 | 100 | 0 | to exp | to exp | 67 |
